# Cure or Curse? Simulation Indicates that Microbes Proliferate under Disinfection Measures in the Space Station

**DOI:** 10.1101/2024.04.16.589799

**Authors:** Shuaishuai Li, Xiaolei Liu, Letian Chen, Hong Liu, Dawei Hu

## Abstract

The long-term stay on the space station requirement poses a knotty issue about the stable hardware and reliable guarantee of astronauts’ health. Despite regular cleaning with wipes, microbes still proliferate in the space station. This study distilled three key factors—low-dose ionizing radiation, dilution and quaternary ammonium compound from the scenario. Species abundance and differential metabolites were applied to test their sole/combined influences on the three-simple-species microbial community succession. Mechanisms were built in mathematical models to generate the structural and behavior similarity. Results showed that LDIR, dilution and QAC, solely or jointly, could contribute to microbial proliferation. The synergy of disturbances might convert them from harmful factors to rewarding ones (e.g. the transformation of QAC into nutrients under LDIR), leading to this “Ecological Surprise”. These results shed light on mechanisms driving microbial community succession in the space station and highlight the need for tailored biocontrol strategies in the specific environment.

## 1. Introduction

Long-term residence on the space station needs stable hardware and guaranteed health status of astronauts (Brooks et al., 2014; Rosado et al., 2010; Sun et al., 2016; Yang et al., 2020), which is threatened by proliferative microbes on the inner surface of the space station (Alekhova et al., 2005). When environmental factors are the same, the species diversity on the space station is much higher than that on the ground (Lang et al., 2017; Novikova et al., 2006), showing the paradox of the plankton (G. E. Hutchinson, 1961). As astronauts and goods are main sources of microbes in this closed environment (Lax et al., 2014; Singh et al., 2018), periodic cleaning is an actual plan for it. When carrying out the cleaning task, astronauts use wipes containing 0.1% quaternary ammonium compounds (QAC) (Singh et al., 2018) to wipe down the internal surface of the cabin under the low-dose ionizing radiation (LDIR) from the universe. However, the removal of microbes seems to be only effective for a short period of time and it is not complete. Microbes always grow back soon after and the community structure varies by time and location (Lang et al., 2017; Singh et al., 2018).

Understanding community succession mechanisms is always one of core targets of ecology, especially microbial ecology(Powell et al., 2015; Stegen et al., 2013; Zhou & Ning, 2017). Disturbance, as a largely independent area of ecology, is the event that influences populations, resources or the physical environment in community succession (Pickett & White, 1985). Removal by wiping, translated into the perspective of microbial community succession, is caused by abiotic environmental disturbance. This operation results in microscopic mass non-specific or specific death, macroscopic physical dilution and chemical disinfection events. Community succession occurs when organisms are disturbed due to their sensitivity to the abiotic environment (Baudoin et al., 2008; Dangles & Chauvet, 2003; Tolkkinen et al., 2013). Although violent environmental degradation might lead to species extinction, some species could be favored under that circumstance because of the interactions necessary for survival in multi-species communities (Abreu et al., 2019, 2020). For example, evenly increasing mortality favors the faster grower, reversing the winner from the slower to faster one in competition (Hastings, 2013; Stewart & Levin, 1973). Nevertheless, current oversimplified models might lose some key information about the succession. Research to date has not yet determined how disturbances would work in resource-explicit models and the influence of selective disturbances.

In this study, three different types of disturbances, LDIR, dilution and QAC, were distilled from the scene of wiping the inner surface of the space station. Experiments with the simple three-species microbial community were carried out to demonstrate species absolute abundance and differential metabolites in the community succession under different combinations and gradients of these disturbances. Results showed that trends of community succession varied among different treatments and all types of disturbances could cause the proliferation by the relief of competition or property transformation of specific chemicals. Afterward, some measures were proposed for microbe control in aerospace. Also, it is hoped that this research will contribute to a deeper understanding of community succession formations and mechanisms under different types of disturbances.

## 2. Materials and methods

### 2.1 Strains, culture and radiative environment simulation

Three laboratory-stored strains, *Bacillus subtilis* (CMCC(B) 63501), *Escherichia coli* (CMCC(B) 44102) and *Pseudomonas aeruginosa* (CMCC(B) 10104) were originally obtained from China Medical Culture Collection. These three species have been reported many times from microbial surveys in the space station, spacecrafts as well as related studies(Crabbé Aurélie et al., 2011; Singh et al., 2018; Urbaniak et al., 2018; Yang et al., 2021; Zea et al., 2016) and it was quite easy to distinguish them by genetic relationship.

To simulate the nutrient-deficient environment on the inner surface of the space station, overnight cultures of above strains were inoculated in the 10ml modified minimal R2A broth (Yang et al., 2021) with a total density of 1×10^6^/ml calculated by direct counting under microscopy (Z. Song et al., 2015). The composition of modified minimal R2A was shown in Table S1. In the mixed culture, the initial ratio of three species was 1:1:1.

The LDIR environmental simulator (Fig. S1-A) was a refitted artificial climate chamber that simulated the LDIR environment in the space station. The modification included three main parts: a rotating wheel for placing cultures, two X-ray tubes for alternatively ionizing the air inside to plasma, and a protective door for isolating the radiation. Two X-ray tubes (XCH201, Dandong Shenbo Electronics Co. Ltd, China) were installed at the inside top of the chamber and run at 20kV, 0.2mA, alternately. To ensure that each culture is exposed to the same radiation dose, six boxes were affixed to the rotary wheel running at a constant angular velocity. These boxes contained plates with LDIR-treated cultures, while control cultures without LDIR treatment were placed outside the radiation-exposed area but in the same climate chamber. The protective door was sealed to maintain a constant temperature of 30°C, and normal pressure during operation. 3D geometric configuration of LDIR environmental simulator was accurately established based on its internal dimensions (Fig. S1-B).

### 2.2 Periodic sampling and successive incubation

During a seven-day experiment cycle, 1ml samples were taken from cultures at the same time on each day. Then the residual parts were differently treated for successive incubation by distinct groups in Table S2. Each group contains three replicates. In each 1ml sample, 700μl was centrifuged for untargeted metabolomics test with its supernatant, and two portions of 90μl bacterial fluid were used to depict species abundance curves.

The composition of QAC was shown in Table S3.

### 2.3 Species abundance curve depicted by qPCR

Two portions of 90μl bacterial fluid above were applied to depict curves for *Bacillus subtilis* (Kirchhoff & Cypionka, 2017; Tut et al., 2021; Xie et al., 2016) by direct qPCR (Sup. A) and *Escherichia coli* as well as *Pseudomonas aeruginosa* by PMA-qPCR, separately. For the PMA-treated one, 10μl 500mmol/L PMAxx (Cat. 40069, Biotium Inc., Hayward, CA, USA) was added into the sample under dark conditions for forming the 50mmol/L final concentration. Then the mixed solution was incubated at 20 for 40 min in the dark on a vortex with mild continuous shaking (100rpm) followed by the change of tube. Subsequently, the tube was exposure to a 120W blue LED light at 4 for 30 min (Sup. B).

DNA was subsequently extracted from the solution above (with PMA or not) by direct boiling with glass beads (Li et al., 2023). 0.1g 0.1mm glass beads (Cat. 11079101, BioSpec Products, Inc., Bartlesville, OK, USA) were added into the tube containing 100μl resuspending solution before the heating procedure. After centrifugation, the supernatant is DNA solution.

Primers, their qPCR standard curves, their amplicons, the composition of qPCR reaction buffer, and the qPCR amplification program were the same as our previous study (Li et al., 2023). 2×SYBR Green PCRmix (Cat. SR1110, Solarbio Science & Technology Co. Ltd., Beijing, China) was used to provide PCR Buffer, MgCl_2_, dNTPs, Hs Taq DNA Polymerase, SYBR Green and so on. The reaction was performed using Gentier 96C real-time PCR system (Tianlong Science and Technology Co. Ltd., Xi’an, China). The qPCR amplification program included: pre-denaturation at 95°C for 10 min with a heating rate of 6°C/s; then 40 cycles of denaturation at 95°C for 10 sec with a temperature change rate of 5°C/s, annealing at 60°C for 20 sec with a cooling rate 5°C/s and elongation at 72°C for 30 sec with heating rate 6°C/s.

### 2.4 Untargeted metabolomics test

Metabolites were quantified using ultra-performance liquid chromatography-triple quadrupole time-of-flight mass spectrometry (UPLC-MS) (Waters Corp., Milford, MA, USA). Raw data obtained from UPLC-MS were quantified in Progenesis QI (Waters Corp.). Significantly differential metabolites analysis was performed using BMKCloud (www.biocloud.net).

### 2.5 Mechanical modeling and computer simulation

The kinetic model was established based on generalized Lotka-Volterra (gLV) equations and hypothetical mechanisms of disturbances. Parameters implying attributes in the community were embedded into the model. Afterward, a corresponding simulation model with dde23 solver was established by S-Function of MATLAB/Simulink (MathWorks, Inc., Natick, MA, USA) to confirm the proposed hypothesis through behavior similarity between computer simulation and experimental data.

## 3. Results

This study aimed to demonstrate the influence of three different types of disturbances on microbial community succession. Experiments were taken to depict the species absolute abundance and differential metabolites during the succession. The following mechanical models and computer simulation were proposed to explain mechanisms of phenomena in these experiments.

### 3.1 qPCR results

The overview of all conditions was shown in Fig. 1. As seen in this figure, distinct succession could be obtained by different treatments. Two-way repeated measured ANOVA was applied using R (4.0.2) to calculate p-values to test if curves showed a significant difference between the specific experimental group and the control group in the overall experimental period, where the interaction effect of treatment and time was considered first. If the interaction effect was not significant, the main effect of treatment would be applied, where the p-value was marked as “*p_m_*”.

**Figure 1:**
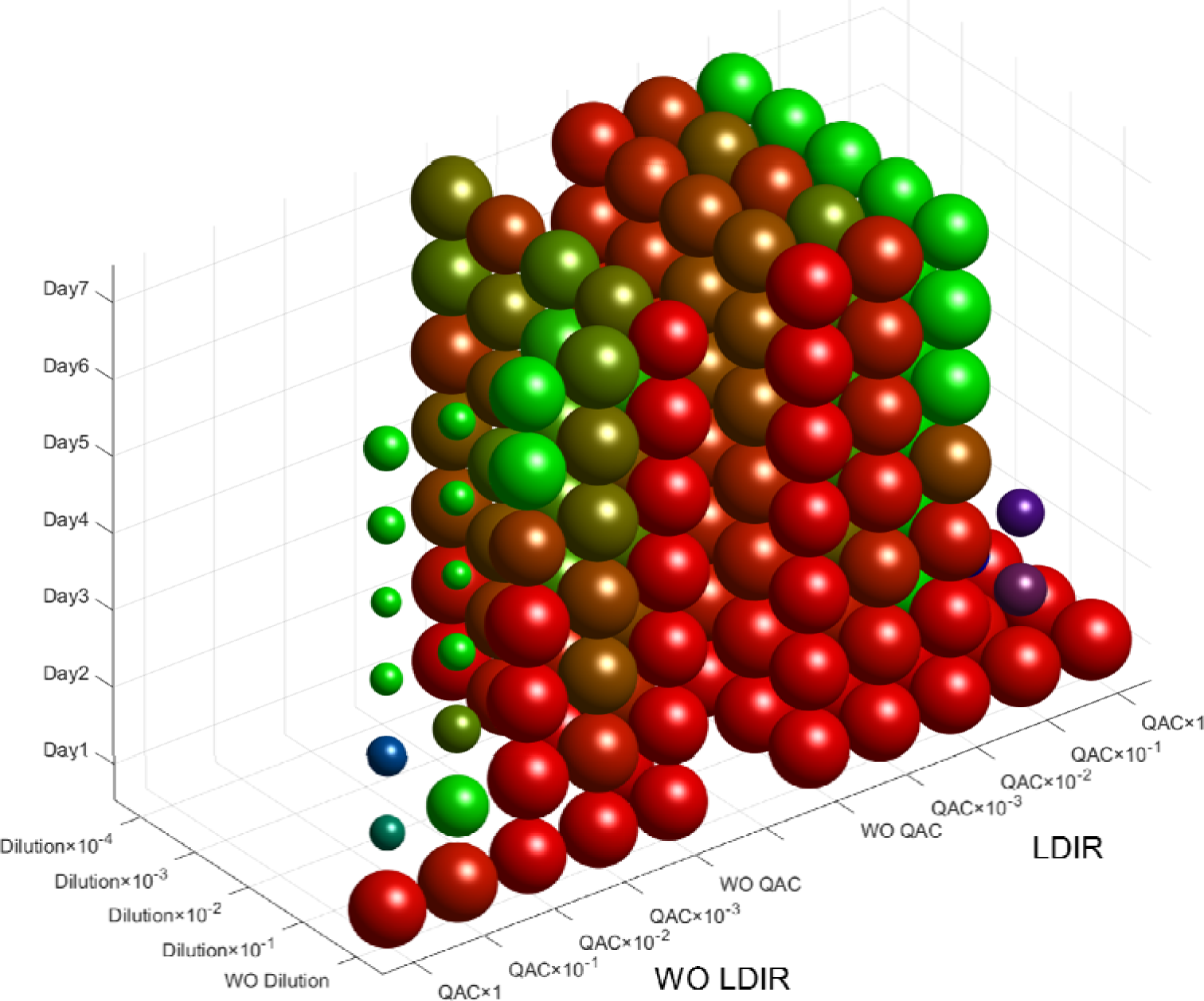
The overview of the community succession under all treatments. The x-axis represents the QAC concentration (e.g. 1 stands for 100%, 10^−1^ stands for 10%), where the left side of axis includes groups without LDIR and the right side includes groups with LDIR; y-axis represents the dilution ratio (e.g. 10^−1^ stands for 10%, 10^−2^ stands for 1%). each position on the x-y plane corresponds to a different treatment. z-axis denotes the number of days, and the seven spheres in each cluster mean seven days of community succession for a treatment group. The radius of the spheres indicates the total biomass in the community (compared to the maximum value and taken as log ten). RGB color shows the percentage of different species (Blue, red and green stand for *Bacillus subtilis*, *Escherichia coli* and *Pseudomonas aeruginosa*, respectively). WO means “without”. The 3D version is uploaded as a .fig file in the supplementary materials.

As shown in Fig. S2, in the control group, the total biomass and the absolute abundance of *E. coli* reached highest points on the sixth day, when the density for both of them was about 1.5 x 10^10^/ml.

When exposed to LDIR (Fig. 2), the microbial community thrived after a period days and still peaked on the 6^th^ day, approximately reaching 3.5 x 10^10^/ml. The of “hibernation”. Compared with the control group, the total biomass rose on 5-7^th^majority was still *E. coli*. The abundance of *B. subtilis* showed the trend of damped periodic oscillations.

**Figure 2:**
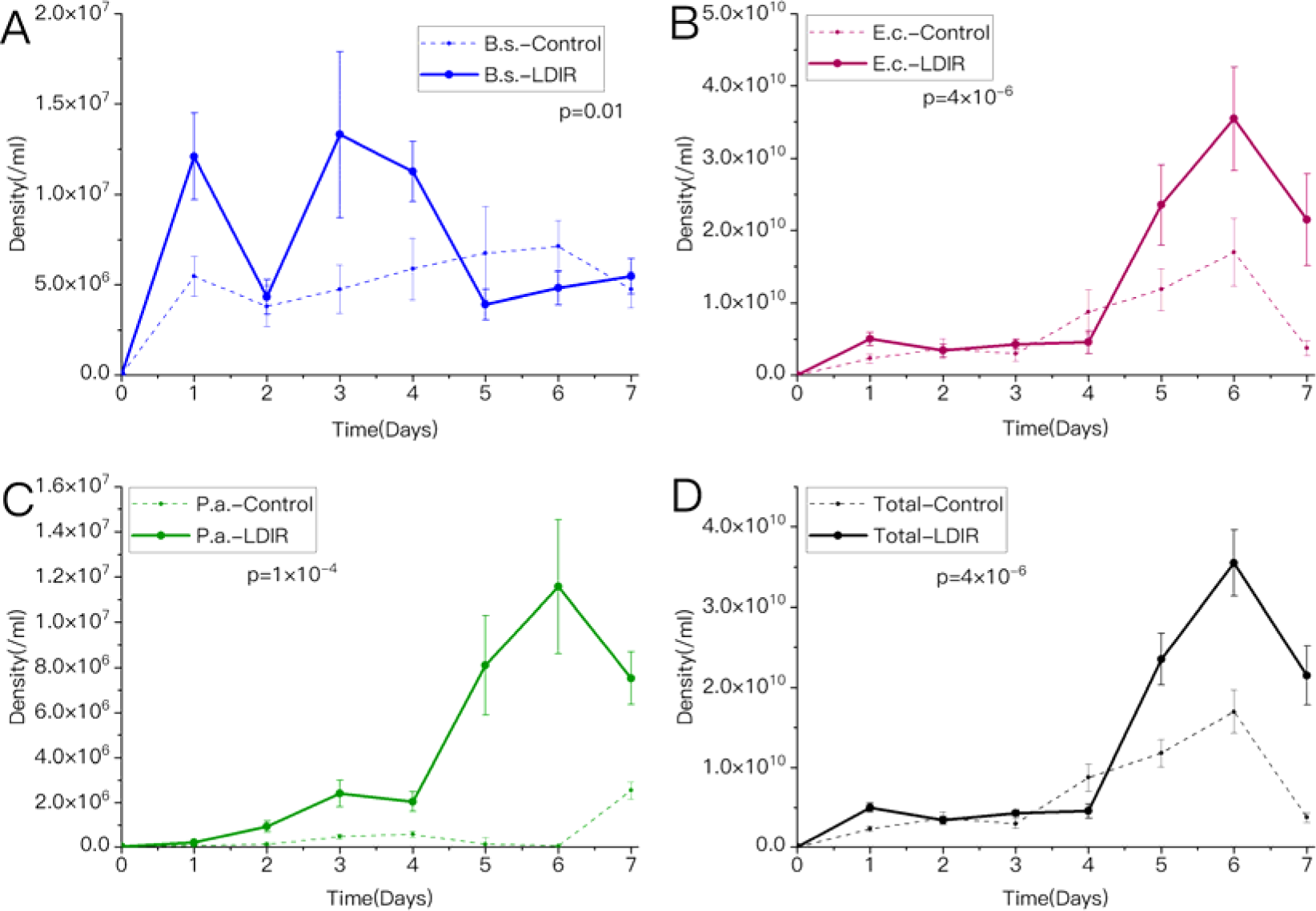
The absolute abundance of three species (A-C) and the total biomass (D under LDIR (solid line) and in the control group (dotted line).

Dilution with gradients from 10% to 0.1‰ per day had no obvious different influence with each other, as shown in Fig. S3 & S4. Taking 1‰ as an example (Fig. 3), compared with the control group, the total biomass and the abundance of *E. coli* abundance of *B. subtilis* significantly fell (*p_m_* = 1 x 10^−5^), while that of decreased on 4-6^th^ days, exhibiting the trend from oscillation to robustness. The *P. aeruginosa* rose dramatically (*p* = 0.004).

**Figure 3:**
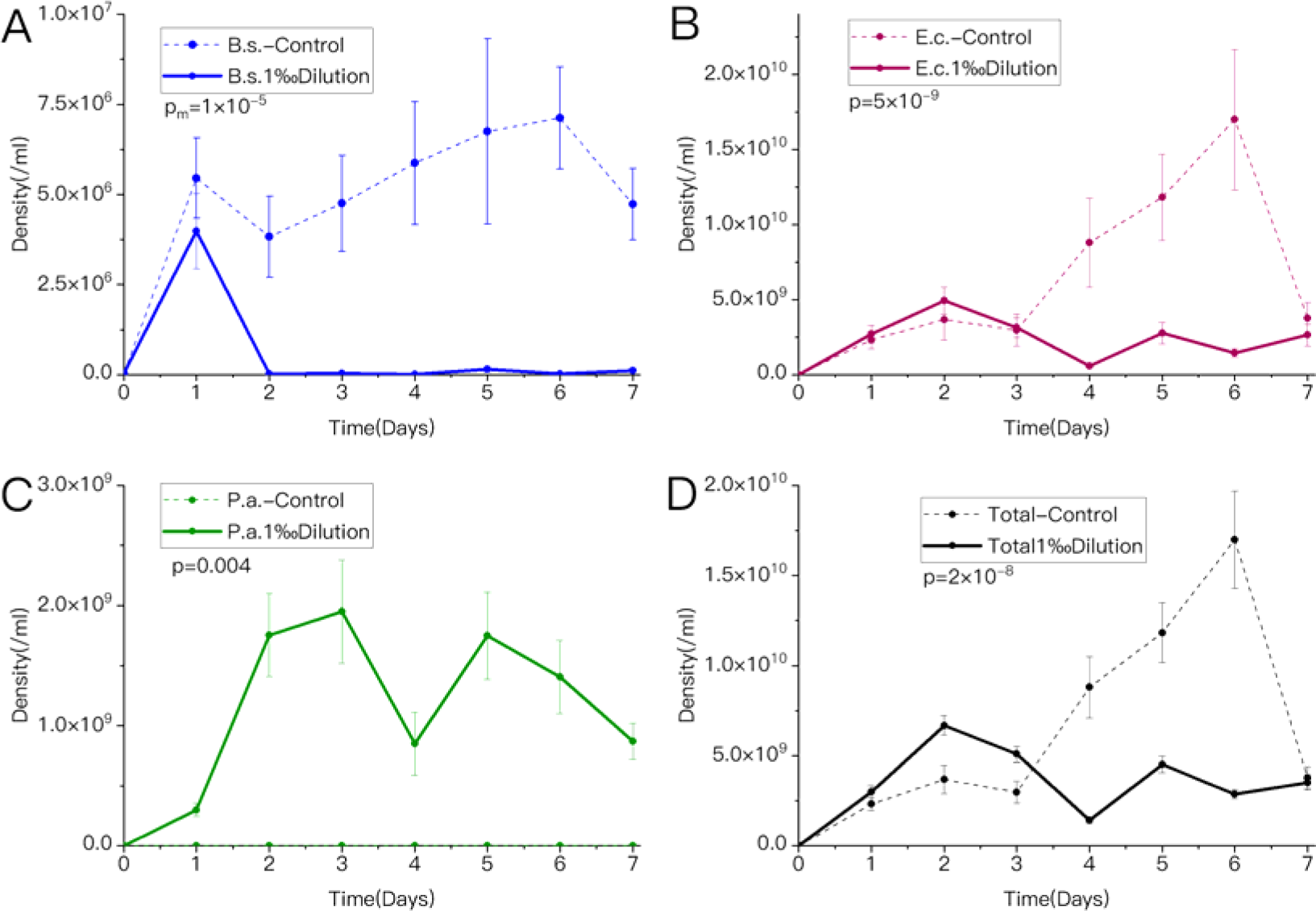
The absolute abundance of three species (A-C) and the total biomass (D) under 1‰ dilution (solid line) and in the control group (dotted line).

Imposing 100% and 10% stock solution of QAC could rapidly bring the total biomass down to below the initial density, when only *P. aeruginosa* could survive below the initial density under this condition (Fig. 1). However, 1% and 1‰ stock solution of QAC did not have the similar impact (Fig. 4). When treated with 1% stock solution of QAC, compared with the control group, the peak of total biomass moved back to 4^th^ day, having no significant difference with the peak of the former. *E. coli* had a great contribution to the total biomass on the 1-4^th^ days, followed by the sharp drop and extinction on the 6th day. Meanwhile, the abundance of *P. aeruginosa* increased from the 5^th^ day and peaked on the 6^th^ day. In the group of 1‰ treatment, the total biomass exceeded the control group on the 7^th^ day before the lag on the 4-6^th^ days, presenting the trend of rise with fluctuations. The abundance of *E. coli* fluctuated continuously, and that of *P. aeruginosa* increased steadily.

**Figure 4:**
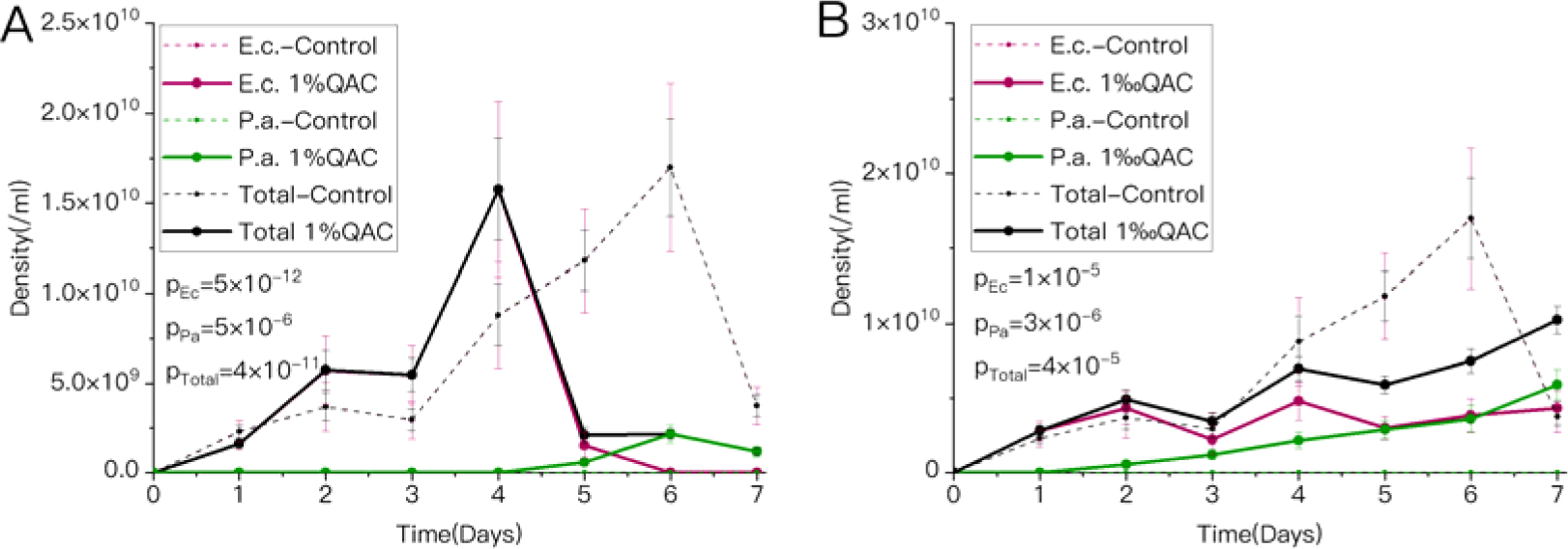
The absolute abundance of *E. coli*, *P. aeruginosa* as well as the total biomass under 1% QAC (A, solid line) and 1‰ QAC (B, solid line) and in the control group (dotted line). *B. subtilis* has a very low abundance, so it is not shown in the figure; the same as below.

When diluted with a 1‰ ratio per day, the total biomass (*p_m_* = 4 x 10^−4^) and the abundance of *E. coli* (*p_m_* = 4 x 10^−4^) significantly rose in the whole period of the LDIR group, compared with the group without LDIR (Fig. S5).

When exposed under LDIR, compared with the group without QAC, the total biomass of the 1% stock solution of QAC group increased on the 2-4^th^ days but decreased on the 5-7^th^ days, in which the abundance of *E. coli* went extinct quickly on the 5^th^ day and that of *P. aeruginosa* peaked on 6^th^ day (Fig. S6-A). The total biomass of the 1‰ stock solution of QAC group rose in fluctuations, exceeding the group without QAC on the 4^th^ and 7^th^ days but lagged on the 5-6^th^ days, when the latter group was experiencing proliferation (Fig. S6-B). The majority was still *E. coli* but the magnitude of *P. aeruginosa* also improved.

When treated with 1% stock solution of QAC, the total biomass of the LDIR group went up in the whole period compared with the group without LDIR except on the 4^th^ day, when the latter group peaked. The abundance of *E. coli* increased on 1-3^rd^ days, then dropped on 4^th^ day but this species still survived later. The abundance of When treated with 1‰ stock solution of QAC, the total biomass (*p* = 0.003) and the *P. aeruginosa* rose during the whole period and still peaked on the 6^th^ day (Fig. S7-A). abundance of *E. coli* (*p =* 0.02) of the LDIR group dramatically rocketed with growing gaps in the whole period compared with the group without LDIR (Fig. S7-B). The abundance of *P. aeruginosa* shrank a little but the difference was not significant (*p >* 0.05).

When LDIR, 1‰ dilution and 1% stock solution of QAC were imposed, compared with the control group (Fig. S8-A), *E. coli* went extinct quickly on the 2^nd^ day. However, the abundance of *P. aeruginosa* substantially increased (*p =* 3 x 10^−6^). When LDIR, 1‰ dilution and 1‰ stock solution of QAC were imposed, compared with the control group (Fig. S8-B), the total biomass increased on 1-4^th^ and 7^th^ days, showing a trend of fluctuation. The abundance of *P. aeruginosa* markedly increased (*p =* 3 x 10^−4^).

### 3.2 Untargeted metabolomics test results

Samples involved in metabolite analysis were divided into 24 series according to biological repeat, and the symbol corresponding to the experimental condition of each group was shown in Table 1. The quality control, OPLS-DA as well as its validity test, OPLS-DA permutation test and screening of metabolites were done before analysis (Sup. C).

**Table 1.** The experimental conditions of distinct metabolomics test series.

| Series | LDIR | Dilution | QAC | Day |
| --- | --- | --- | --- | --- |
| A | N | N | N | 1 |
| B | N | N | N | 4 |
| C | N | N | N | 5 |
| D | N | N | N | 6 |
| E | N | N | N | 7 |
| F | N | 1‰ | N | 4 |
| G | N | N | 1% | 4 |
| H | N | N | 1% | 5 |
| I | N | N | 1% | 6 |
| J | N | N | 1‰ | 4 |
| K | N | N | 1‰ | 7 |
| L | Y | N | N | 1 |
| M | Y | N | N | 4 |
| N | Y | N | N | 5 |
| O | Y | N | N | 6 |
| P | Y | N | N | 7 |
| Q | Y | 1‰ | N | 4 |
| R | Y | N | 1% | 4 |
| S | Y | N | 1% | 5 |
| T | Y | N | 1% | 6 |
| U | Y | 1‰ | 1% | 4 |
| V | Y | N | 1‰ | 4 |
| W | Y | N | 1‰ | 7 |
| X | Y | 1‰ | 1‰ | 4 |

Compared with the control group, the concentration of p-coumaroyl agmatine in the group exposed under LDIR between 4-6^th^ days dropped significantly (*p <* 0.05, Series B&M, C&N, D&O. Fig. S9). On 1^st^ and 4-6^th^ days, most of metabolic pathways were inhibited, including polysaccharide metabolism, amino acid metabolism, vitamin metabolism, xenobiotic degradation and metabolism, as well as antibiotic biosynthesis (A&L, B&M, C&N, D&O; Fig. S10).

Compared with the control group, in the group diluted with 1‰ ratio per day, N-acetylmannosamine was enriched on the 4^th^ day (B&F), which was markedly correlated with the rise of the abundance of *P. aeruginosa* (*p* < 0.01, Fig. S11). Pathways inhibited dominated on 4^th^ day, including nucleotide metabolism, sugar degradation, antibiotic synthesis, and aminobenzoate degradation (Fig. S12).

When treated with 1% stock solution of QAC, N-acetylmannosamine was more abundant on 4^th^ day compared with the control group (B&G), which was also correlated with the rise of the abundance of *E. coli* (*p* <0.05, Fig. S13). Meanwhile, staurosporine and tetracycline biosynthesis were active. When treated with 1‰ stock solution of QAC, echitovenine was rarer on the 7^th^ day compared with the control group (E&K), which was also correlated with the increase of the abundance of *P. aeruginosa* (*p* <0.01, Fig. S14). On the same day, aminobenzoate degradation as well as biosynthesis of alkaloids (derived from histidine and purine) were inhibited (Fig. S15).

When both LDIR and 1% stock solution of QAC were imposed, compared with the group under 1% stock solution of QAC, alkaloid biosynthesis was inhibited (Fig. S16), and some nitrogenous long-chain organic matters were enriched on 5-6^th^ days ethanolamide, ranging from 1 x 10^10^ ∼ 2 x 10^12^ times; compared with the control (H&S, I&T), including tetradecenoyl carnitine and R-palmitoyl-(2-methyl) group, cytochrome P450 metabolism was active on 5^th^ day (C&S). When LDIR, 1‰ N-[(2R)-2-hydroxypropyl] hexadecanamide was 2 x 10^12^ times higher than the dilution and 1% stock solution of QAC were imposed together, the concentration of control group on the 4^th^ day (B&W). However, when the QAC concentration was changed to 1‰ of the stock solution (B&V, B&X, E&W, J&V, K&W), the phenomenon of enrichment of nitrogenous long-chain organic matters disappeared; alternatively, cyanoamino acid metabolism, related to nitrogen metabolism (Ding et al., 2022), was active. The concentration of albaflavenone sharply diminished (*p* < 0.01, Fig. S17) on 4^th^ day in the group with both LDIR and 1‰ stock solution of QAC, compared with the control group (B&V). Aminobenzoate degradation on the 4^th^day and antibiotic synthesis on the 7^th^ day (K&W. Fig. S18) were restrained in the group with both LDIR and 1‰ stock solution of QAC, compared with 1‰ stock solution of QAC group. Under LDIR, when comparing 1% and 1‰ stock solution of QAC groups, eicosanoids metabolism was active in the higher concentration one on the 4^th^ day (R&V. Fig. S19).

### 3.3 Mechanical modeling and simulation results

#### 3.3.1 State variables

The mechanical model was composed of six state variables in four types: *B. subtilis* biomass (*x_1_*), *E. coli* biomass (*x_2_*), *P.aeruginosa b*iomass (*x_3_*); amount of substrate (*S*), amount of QAC (*Q*), and amount of QAC degradation products (*P*).

#### 3.3.2 Rate equations

1. growth rate The growth rate without competition of species could be expressed as:

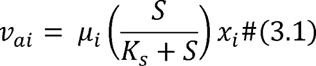

where *μ_i_*was the specific growth rate of species *i*, and *K_s_* denotes substrate-dependent half-saturation constant for species *i*.
2. interspecific competition When there was no joint treatment of LDIR and QAC, the interspecific competition rate against species *i* could be written as follows:

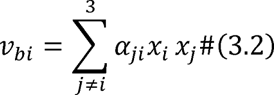

where *α_ji_* denotes the competition inhibition coefficient from species *j* to species *i*. When the microbial community was treated with LDIR or QAC, the interspecific competition rate against species *i* could be written as follows:

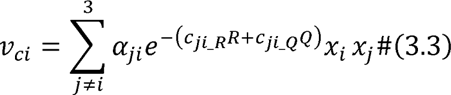

Where *I* was a constant, representing the LDIR coefficient; *e* was the Euler number; *c_ji_R_* and *c_ji_Q_* expressed the interspecies cross-feeding response coefficient between species *j* and *i* species caused by LDIR and QAC, respectively.
3. intraspecific competition When there was no joint treatment of LDIR and QAC, the intraspecific competition rate of species *i* could be written as follows:

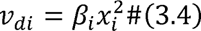

where β*_i_* denoted the intraspecific competition inhibition coefficient of species *i*. When the microbial community was treated with LDIR or QAC, the intraspecific competition rate of species *i* could be written as follows:

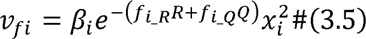

where, *f_i_R_* and *f_i_Q_* denoted the intraspecies cross-feeding response coefficient of species *i* caused by LDIR and QAC, respectively.
4. metabolism The metabolism rate of species *i* could be expressed as:

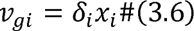

where δ*_i_* was the metabolism coefficient of species *i*.
5. QAC disinfection

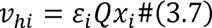

where ε*_i_* was the QAC disinfection rate on species *i*.
6. QAC degradation

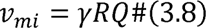

where γ was the first-order kinetic coefficient of QAC degradation rate.
7. generation of QAC degradation products

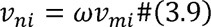

where ω was the calibration coefficient.
8. consumption of substrate

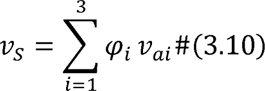

where φ*_i_* was the substrate consumption coefficient during the growth of species *i*.
9. transient influence of Dilution and QAC
  a. only dilution

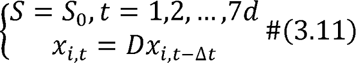

where *S_0_* was the original substrate concentration of the R2A culture; *x_i,t_* was the abundance of species *i* at *t* moment; Δ*t* was a very short time interval; *D* was the proportion of the culture just before dilution to the total volume.
  b. only QAC

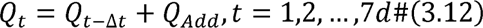

where *Q_Add_* denoted the QAC added.
  c. both dilution and QAC

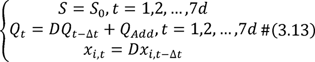

#### 3.3.3 The overall mechanical model

Based on Eq(3.1)∼(3.13), the overall mechanical model, i.e. a state space model with six first-order nonlinear kinetic equations, could be formulated in a top-down manner as follows:

1. When there was no joint treatment of LDIR and QAC, the population dynamics equation of species *i* was:

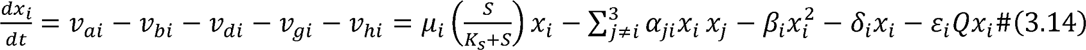 When there was joint treatment of LDIR and QAC, it was:

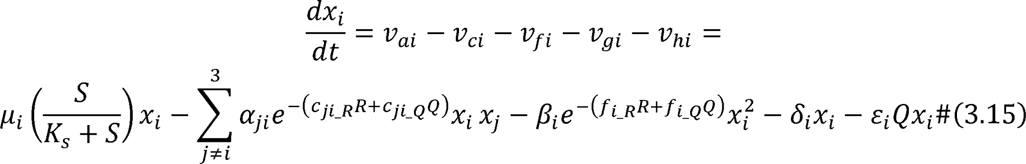
2. The QAC degradation dynamics equation was:

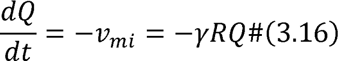
3. The generation of QAC degradation production dynamics equation was:

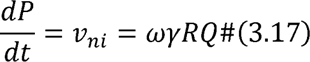
4. The substrate change dynamics equation was:

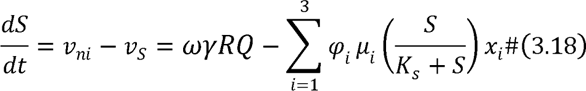

#### 3.3.4 Computer simulation

The code of the mechanical model above was written by S-Function in MatLab/Simulink(R2019b, 9.7.0.1190202). Simulation experiments were implemented for simulating the control group and treatment groups of LDIR, dilution, QAC as well as their combinations, as shown in Fig. S20. The simulation results showed that the mechanical model had considerable behavior similarity with experiments, while the mathematical expression of the mechanical model matched the growth process of microbes, indicating the structure similarity between them. Therefore, the mechanical model was considered valid.

## 4. Discussion

In this study, LDIR, dilution and QAC were extracted from the scenario as treatments, and the microbial community succession under these uniformed different treatments was observed by measuring species abundance and metabolites. Mechanical models were built to explain experiments, with which the computer simulation showed structural and behavioral similarity. The overall mechanisms were shown in Fig. 5.

**Figure 5:**
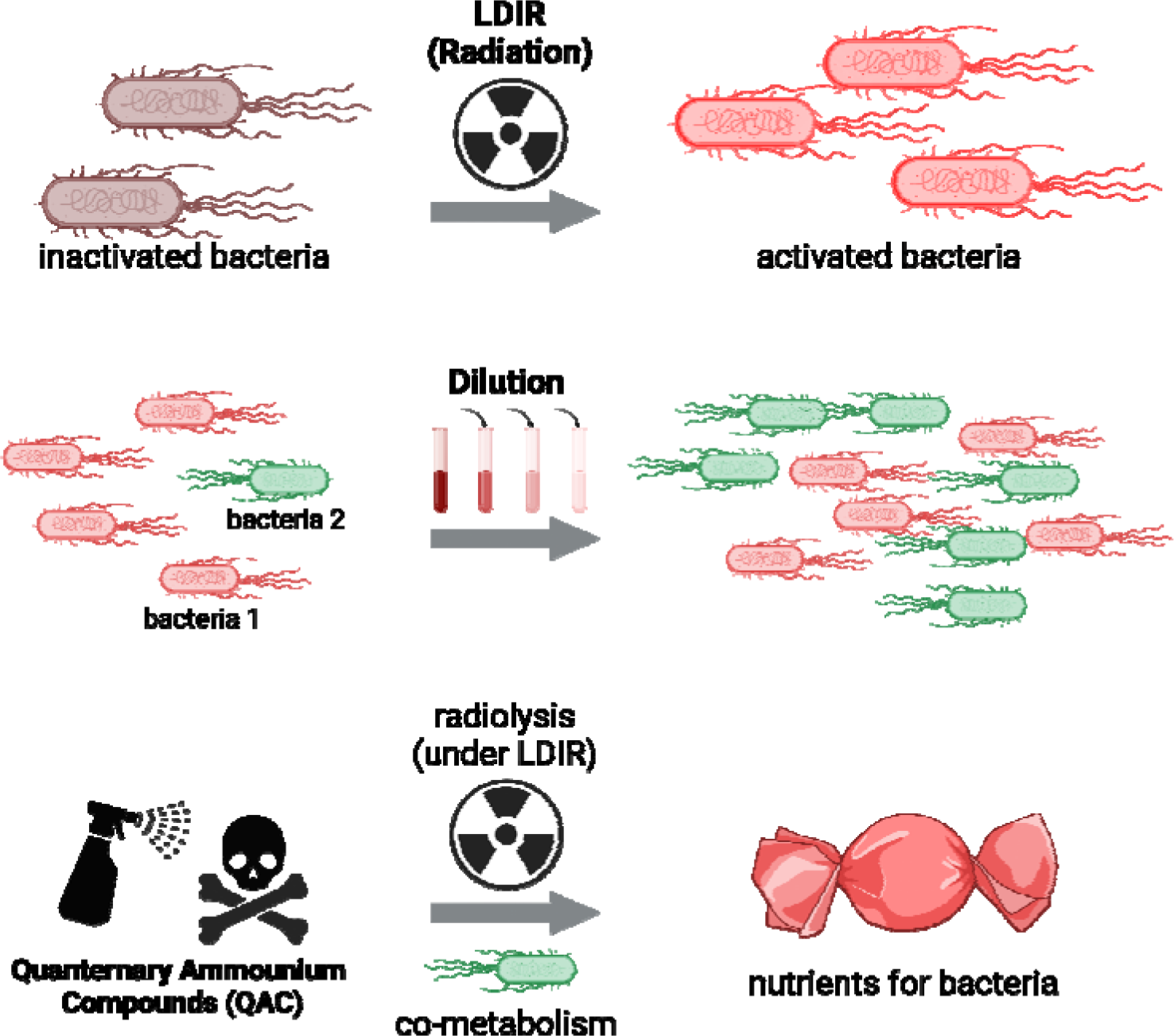
The overall mechanisms of LDIR, dilution and QAC on microbial community succession. LDIR activates all species’ growth and reproduction. Dilution allows specific species to be selectively released from suppression. Low-concentration QAC undergoes radiolysis under LDIR and co-metabolism by bacteria, thus changing from bactericide with harmful effects to nutrients. This figure is created with BioRender.com.

At the later stage, LDIR improved the total biomass, every species abundance and their peaks, when the Hydra Effect happened. This phenomenon was shown in both with/without LDIR comparison and with/without LDIR comparison under dilution (Fig. 2 & S5). P-coumaroylagmatine could thicken cell walls, limit bacterial movement and improve plant defense against pathogens(Häusler et al., 2014; Macoy et al., 2015), thus its decrease on the 4-6^th^ days (B&M, C&N, D&O) indicated that the physiological activity was enhanced under LDIR. The inhibited antibiotic synthesis pathway (B&M) suggested that external pressure was reduced and interspecific competition was diminished. Therefore, the Hydra Effect occurred under LDIR. This effect was more significant as the main effect in the comparison of the group under LDIR and 1‰ dilution and only 1‰ dilution (Fig. S5). Except in *P. aeruginosa*, dilution reshuffled all other components everyday and undermined the interaction effect of treatment and time, where the enhancement effect of LDIR emerged.

Dilution, by removing a certain part of biomass during serial transfers, was a non-specific and highly controllable disturbance way, where the transient removal did not depend on response traits in the microbial community (Gibbons et al., 2016). However, under non-selective dilution, *P. aeruginosa* was selectively released from competitive exclusion. Aminobenzoate was the signal of xenobiotic degradation (Patil et al., 2020; Wang et al., 2020), thus its degradation inhibition on 4^th^ day (B&F) suggested that organisms tended not to consume energy storage material (Smitha et al., 2017) and the competition in the community was mitigated. It could be inferred that many niches would be released without distinction under dilution, when *P. aeruginosa* could reproduce based on its merits beyond others’ (Abreu et al., 2019) and occupy these niches. This release happened only under dilution with all gradients; when both dilution and LDIR were applied, *E.coli* took the majority again (Fig. S21-A).

QAC stood for specific disturbances, causing non-random removal according to the microbe’s response traits(Banitz et al., 2020; Calderón et al., 2018; Jurburg et al., 2017; Srivastava & Vellend, 2005). Its effect also counted on disturbance intensities and frequencies. Different QAC concentrations had various influences on the microbial community, from elimination to promotion. The Hydra Effect even happened under 1% and 1‰ stock solution of QAC, where simple competition mitigation could not explain. Polymers of N-acetylmannosamine played a protective role in *E. coli* (Sellner et al., 2021), thus their respective increments with dramatical correlation told the reason why the species abundance of *E. coli* peaked in advance on the 4^th^ day under 1% stock solution (B&G). Besides, the active biosynthetic pathways of antibiotics and cytokinins on the same day corroborated that microbes combated adversity by more cell division and the total biomass also peaked. Echitovenine was a monoterpene alkaloid with antimicrobial activity (Mohammed et al., 2021), whose attenuation could interpret the growth of *P. aeruginosa* on the 7^th^ day under 1‰ stock solution (E&K), given their notable correlation. QAC would periodically selectively remove a portion of individuals in the community, leading the way of community succession to the direction of dominance by species that can tolerate the environment, which could be pronounced in comparisons across gradients (Fig. S21-B&C). With low-concentration QAC treatment, the community had adapted to the environment and utilized it. QAC, was a series of long-chain organics that could be used by microbes, especially in microbial co-metabolism by *Pseudomonas* (Harris & Knowles, 1983; Leadbetter & Foster, 1960; Zhang et al., 2015), which was widely and systemically utilized in treating wastewater (biological nitrogen removal). On the oligotrophic internal surface of the space station, carbon sources were mostly wiped down, when co-metabolism might allow microbes to survive with remaining nitrogen sources. Meanwhile, the inhibition of aminobenzoate degradation pathway also flanked the corresponding decrease of the consumption of energy storage in the community.

The upsurge of some nitrogenous long-chain organic matters under both LDIR and QAC, compared with only QAC or the control group, strongly implies that QAC may undergo radiolysis under LDIR. Several studies (Baidak & LaVerne, 2010; Dhiman & LaVerne, 2013; Nogami et al., 1996; Pillay, 1986) implied that radiolysis would happen on substrates containing QAC, especially under water acceleration. 1% stock solution of QAC is substantially transformed from the toxic bactericide to organic nitrogenous long-chain nutrients, including tetradecenoyl carnitine, R-palmitoyl-(2-methyl) ethanolamide and N-[(2R)-2-hydroxypropyl] hexadecanamide with significant differences between Series B&W, H&S, I&T, separately. This might accelerate the microbial co-metabolism process. Cytochrome P450-related enzymes prefer long-chain fatty acids with 15-16 carbon atoms (Lewis & Wiseman, 2005), thus its activation might be related to added long-chain QAC utilization. Inhibited alkaloid biosynthesis (H&S, I&T) and ameliorated cytochrome P450 metabolism (C&S) also shows that the community could adapt to and consume QAC under LDIR. Nevertheless, when the QAC concentration is 1‰, significant differences are no longer detected for organic nitrogenous long-chain nutrients (B&V, B&X, E&W, J&V, K&W), suggesting that the community can entirely co-metabolize QAC in this situation. The activation of cyanoamino acid metabolism (B&X), inhibition of aminobenzoate degradation (J&V) as well as inhibition of antibiotic synthesis (K&W) also reconfirm the speculation of utilization of exogenous QAC as the source of nitrogen and less pressure of the community caused by LDIR. Comparing different concentration of QAC under LDIR (V&R), the active eicosanoids metabolism in the higher concentration one showed that microbes might transform QAC into long-chain fatty acid substances for consuming.

The proliferative microbes on the inner surface of the space station (Alekhova et al., 2005) have been threatening its normal operation, which is a serious problem and could be shifted to the issue of the effect of disturbance on community succession. Disturbance is acknowledged as a key factor of biodiversity via affecting eco-processes (Banitz et al., 2020). However, mechanisms behind the observed community diversity under disturbance need to be further clarified (Fox, 2013; Shea et al., 2004). First, few quantitative studies on disturbances have been reported, while more relative studies focus on the qualitative aspect — this is due to both the difficulty in defining the attribute of frequency, intensity, as well as extent (Shade et al., 2012), and the wide variety of their characteristics and ecological impacts (Fraterrigo & Rusak, 2008). Second, few studies are explicitly based on the time course of microbial community composition under disturbance; instead, many solely studies are on the sensitivity of composition at a given moment (Allison & Martiny, 2008), which is attributed to the inertial thinking in community ecology and research tools in microbiology. Our previous study noticed the benefit of sole LDIR for the microbial community (Yang et al., 2021). Abreu et al. (Abreu et al., 2019, 2020) expanded the effect of dilution on multispecies community composition, providing an example for quantifying disturbances with the time course. Nonetheless, in their research, when distinguishing disturbances by pulsed versus sustained, pulsed ones like dilution are not applicable for being directly integrated into the typical LV equation. Finally, studies with similar gradient-based methods on combined multiple factors with synergy, are still at an early stage. Most studies still concentrate on a single factor, and only a few have noticed the combined influence of multiple factors (Banitz et al., 2020; Zeng et al., 2020). In contrast, a large number of phenomena in nature are caused by the complex interaction of the latter condition (Elmqvist et al., 2003; Martínez-Ramos et al., 2016), including “ecological surprises” such as drastic shifts in community compositions and biodiversity (Banitz et al., 2020; Filbee Dexter et al., 2017; Newman, 2019).

In this study, “ecological surprise” occurred when human expectations deviated from the observed community succession behavior (Doak et al., 2008; Filbee Dexter et al., 2017). And these unexpected dynamics arose from human management and disturbances (Doak et al., 2008). Dilution (wiping) and QAC were recognized as efficient measures to take control of microbes (Sattar et al., 2015; Zhang et al., 2015) and applied in the space station (Singh et al., 2018). However, given the results above, the synergy emerged between LDIR and QAC, thus low-concentration QAC was transformed from the bactericide to nitrogen-rich long-chain nutrients. Selectively released *P. aeruginosa* under suppression after dilution would take these nutrients to thrive, changing the community structure and dominant species. That could be a possible reason for the phenomenon in the space station, and also gave a potential mechanism for “ecological surprise” under synergistical disturbances. It reminded us that tailored biocontrol strategies ought to be considered for the specific environment. Sometimes, the control measures were taken for granted that they were “disturbances” affecting targets without the consideration of their own properties and changes. It cannot be neglected that general strategies might not function under certain conditions.

Given the current microbial prevention strategies have some disadvantages, future improvements can be adopted. Concerning the low concentration of QAC, except for co-metabolism and radiolysis of QAC mentioned above, pre-package of wipes can also cause it by both reduced antimicrobial activity of QAC with a long-term soaking (Bloß et al., 2010, 2010; Slopek et al., 2011) and the chemical combination with the wipe fibre (Dharan et al., 1999; X. Song et al., 2019). Target surface properties also influence the disinfection (X. Song et al., 2019). The more organic contamination on (dirtier) the surface, the lower the disinfection effect (Williams et al., 2007). Moreover, radiolysis could happen just after the spacecraft entered the outer space. In terms of dilution, its rate was limited on the order of 1% even using two wipes successively (Gold & Hitchins, 2013), which is not enough given results in this study. Furthermore, “wiping” itself could lead to cross-contamination of microorganisms in different locations by mechanical stimulation (Ramm et al., 2015; Siani et al., 2011; Williams et al., 2007). To attenuate these effects, several measures could be considered, such as augmenting local wiping time, wiping with mechanical equipment to restraint power variation (Sattar et al., 2015; Williams et al., 2007), special antibacterial treatment on the surface requiring microbial prevention, applying different wipe materials harder to react with QAC, adjusting the formulation of QAC for a lower degradation rate (Pillay, 1986; Zhang et al., 2015), avoiding surface exposure to radiation after wiping.

Despite these promising results, questions still remain. Further work is needed to develop reliable experimental and analytical methods (e.g. based on the individual-based model) (Li et al., 2022) for testing community succession considering the spatial factor. The liquid culture implemented in this study aborted it because of current biotechnology limitations on detecting microbial individuals. Additional research based on more methods in bioinformatics is needed to better understand the successive changes in microbial behaviors under disturbances. This study focused on the level of community succession. Hence, there is abundant room for further progress in determining evolutionary characteristics changes of the species facing disturbances in long-term experiments.

## 5. Conclusion

In this study, the intricate relationship between disturbance and the microbial community succession in the space station was elucidated. The proliferation under LDIR, the selective species release under dilution and complex responses of the community to QAC concentration were observed in experiments and their similarity was tested in modeling. The synergy of combined disturbances, QAC and LDIR, leads to the transformation of them from harmful factors to supportive ones. Therefore, it is necessary to consider synergistic effects and to apply tailored strategies when addressing the microbial proliferation challenge as well as other similar circumstances.

## Supporting information

Figure 2 (3D version)

## Abbreviation

LDIR: low-dose ionizing radiation
QAC: quaternary ammonium compound(s)

## Acknowledgment

We sincerely acknowledge the assistance from Chuanmeng Cai, Dengbo Chen, and Yong Feng in the long and non-stop periodic experiment, and the assistance from Yong Feng for depicting Fig.S1-B.

## Data Availability Statement

The data and code used to produce results of this study are openly available in Zenodo at https://doi.org/10.5281/zenodo.11094948.

## Author Contributions

SL, HL and DH conceptualized and designed the study; SL, XL and LC performed experiments; SL and DH built the model and did the computer simulation; SL wrote the first draft of the manuscript; all authors contributed substantially to revisions.

## Supplementary Materials

**Figure S1:**
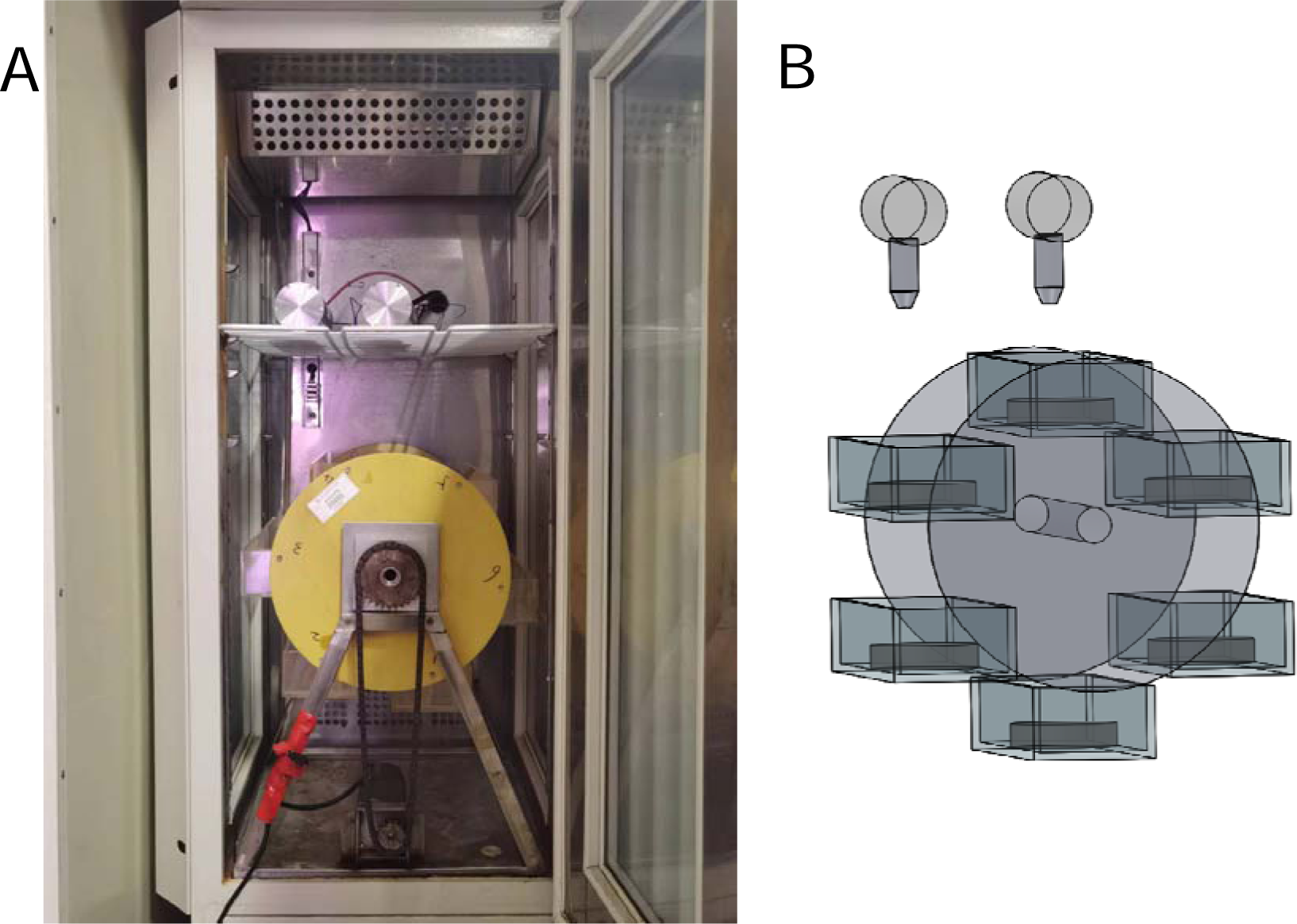
The LDIR environmental simulator (A) and its 3D geometric configuration (B).

**Figure S2:**
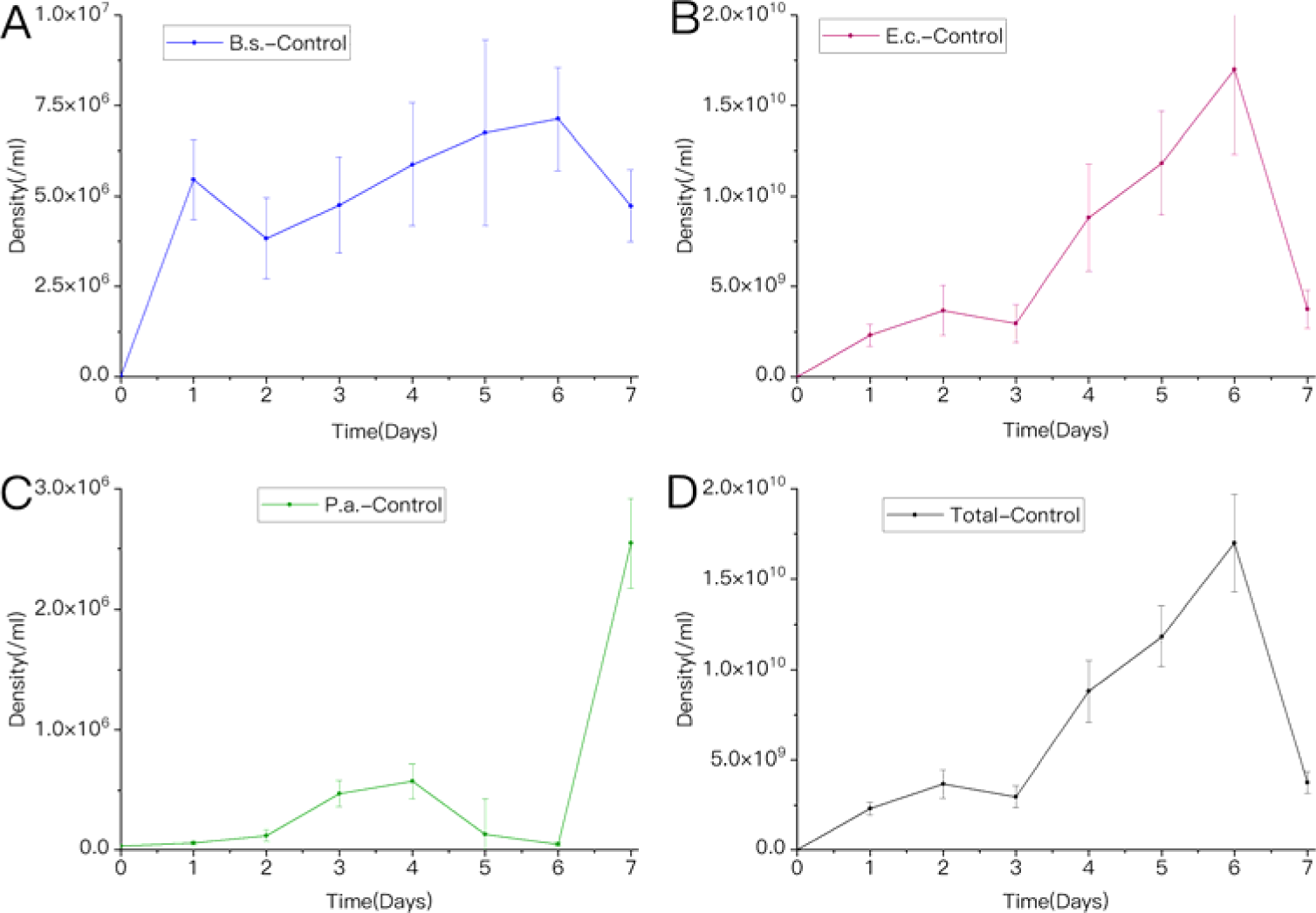
The absolute abundance of three species (A-C) and the total biomass (D) in the control group.

**Figure S3:**
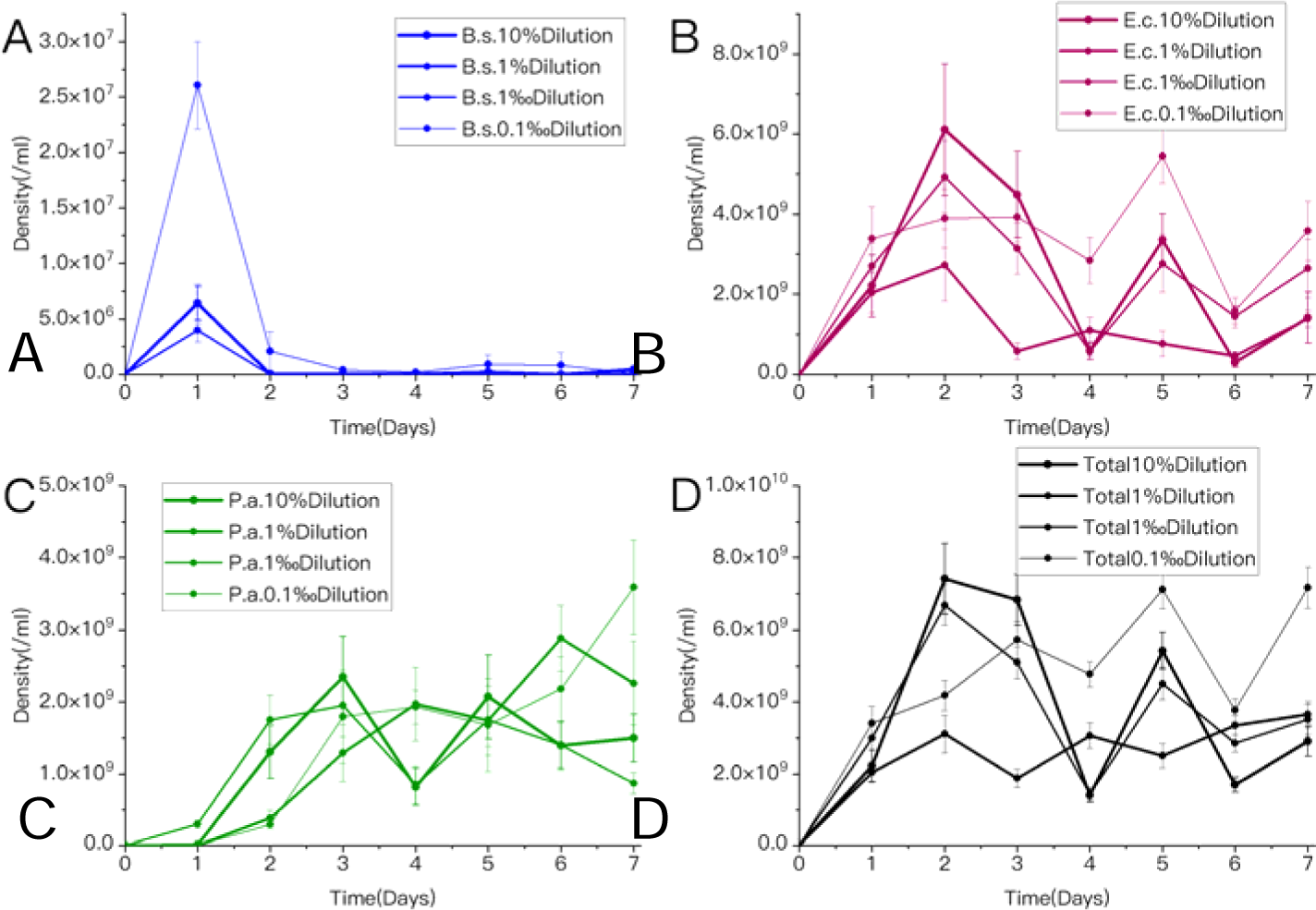
The absolute abundance of three species (A-C) and the total biomass (D) under different dilution ratios. The thinner the line, the more diluted the community in the group is.

**Figure S4:**
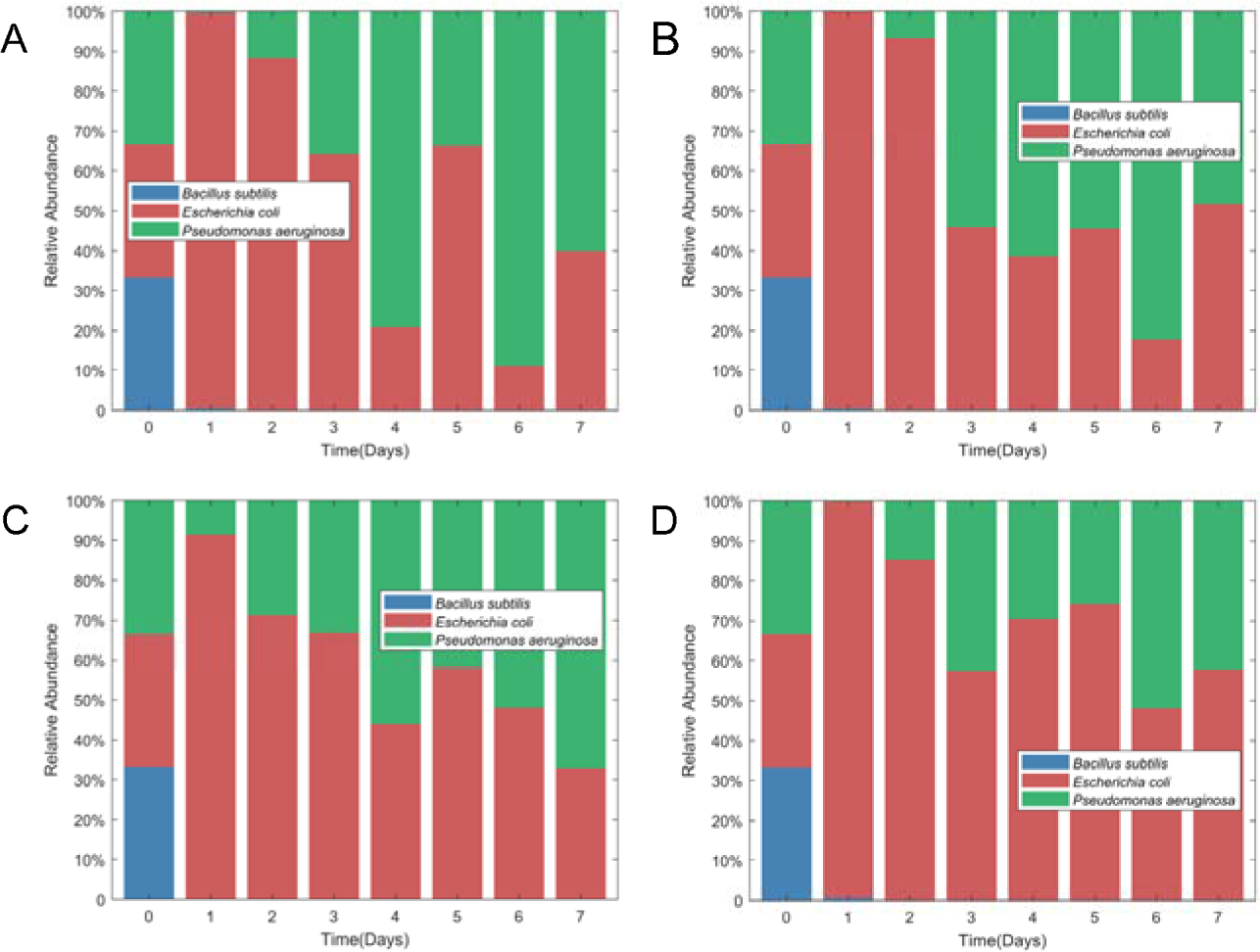
The relative abundance of three species under different dilution ratios. A-D represent 10%, 1%, 1‰ and 0.1‰ dilution ratios, respectively.

**Figure S5:**
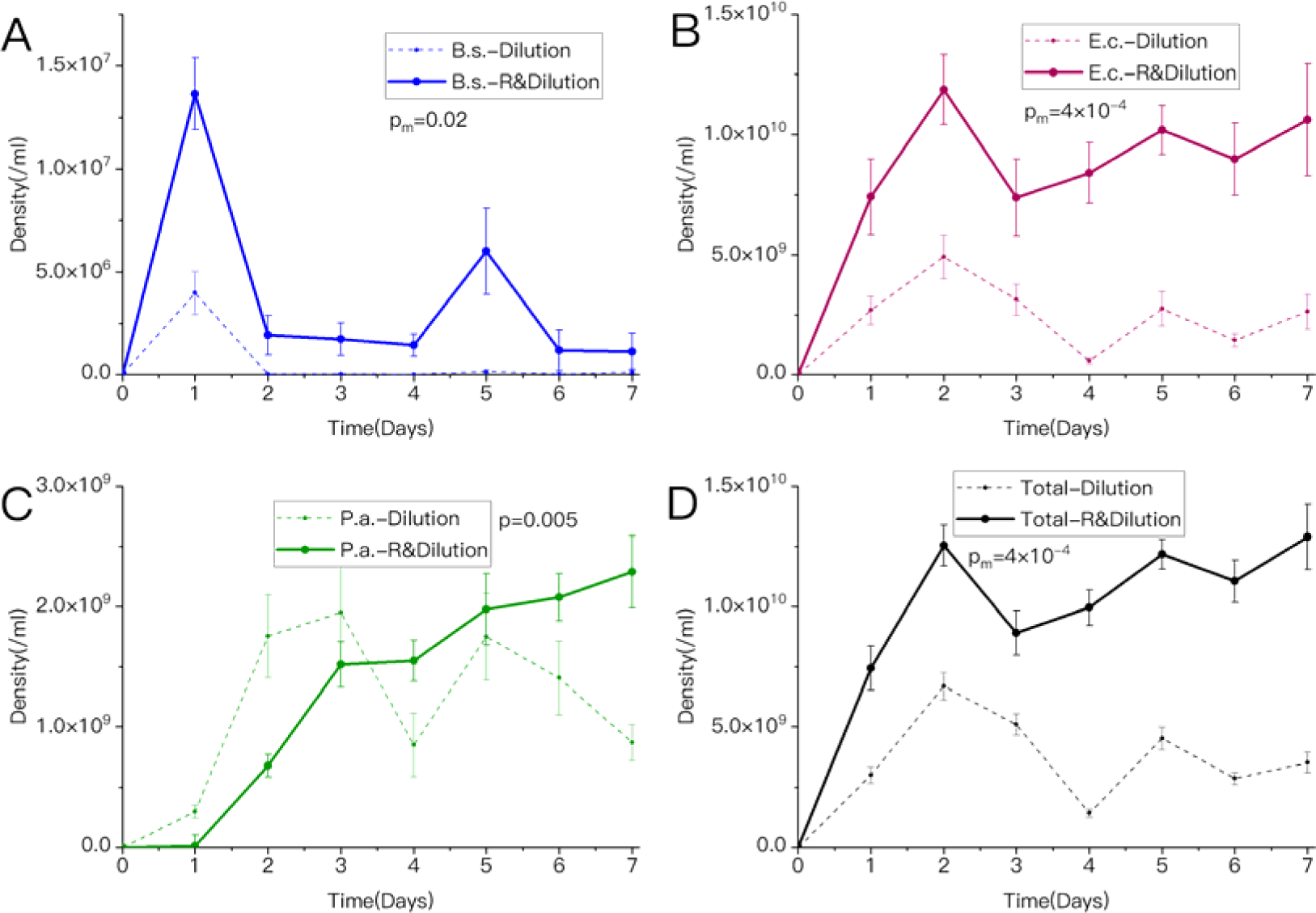
The absolute abundance of three species (A-C) and the total biomass (D) under LDIR and 1‰ dilution (solid line) as well as only 1‰ dilution (dotted line).

**Figure S6:**
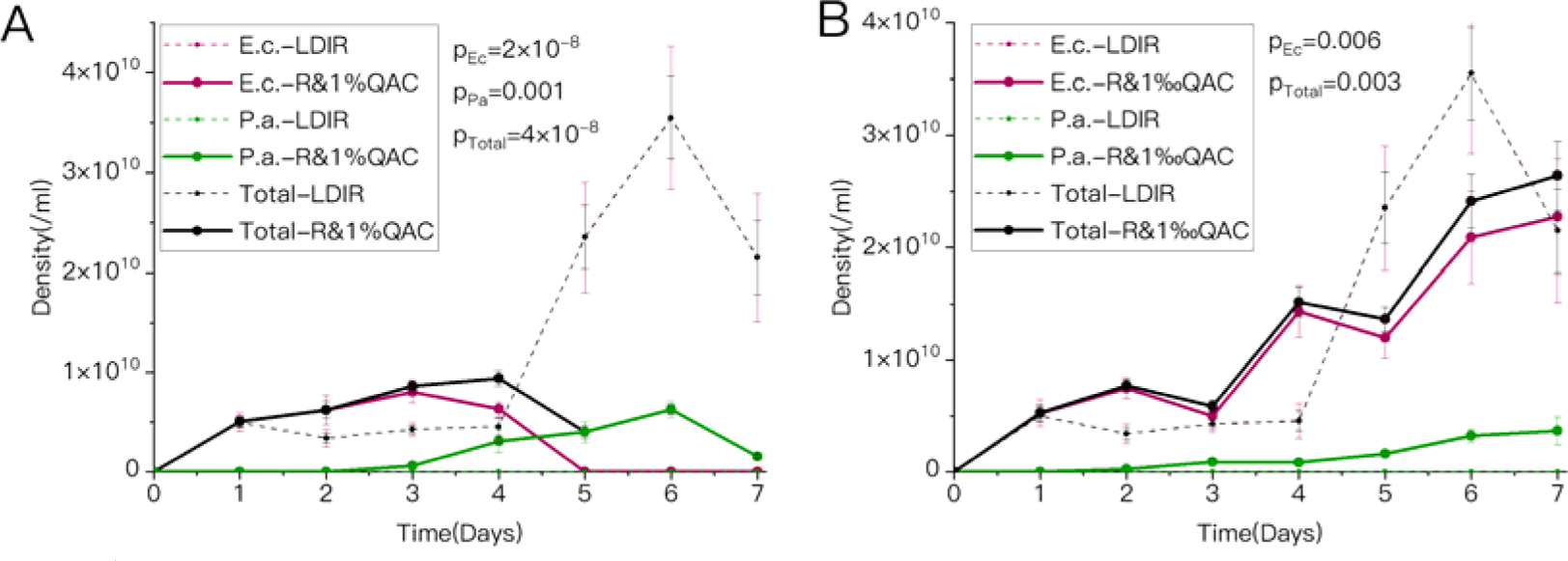
The absolute abundance of *E. coli*, *P. aeruginosa* as well as the total biomass under LDIR and 1% QAC (A, solid line) or LDIR and 1‰ QAC (B, solid line) as well as only LDIR (dotted line).

**Figure S7:**
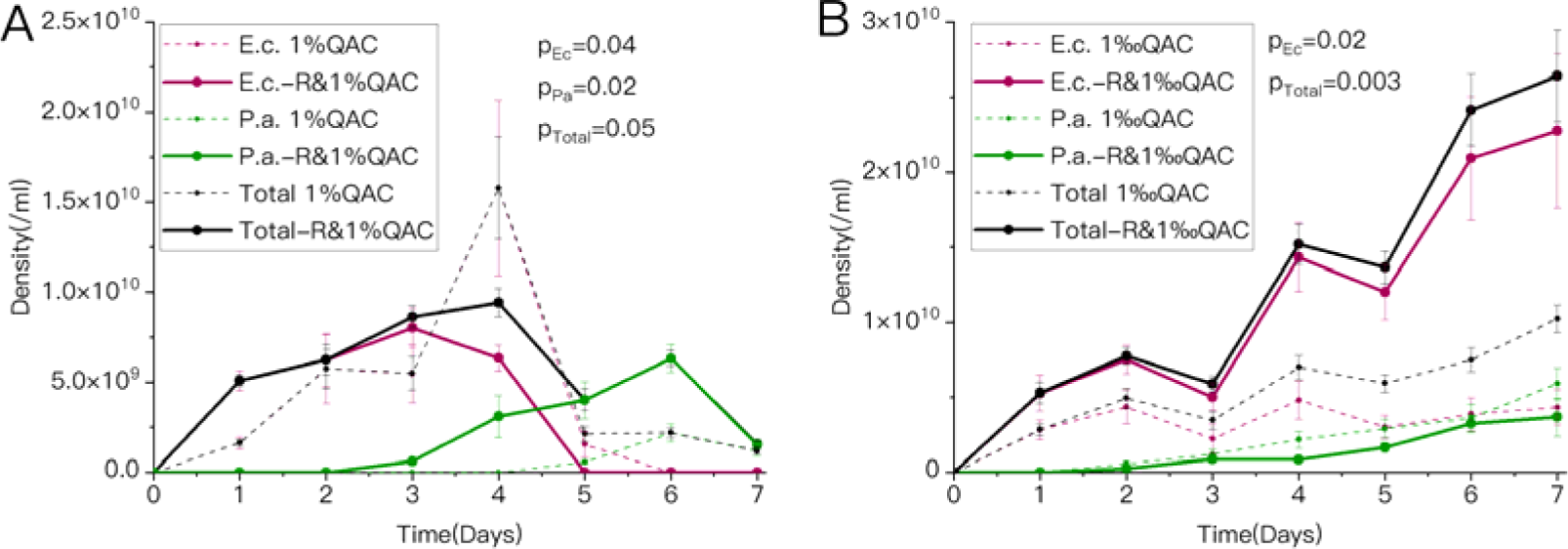
The absolute abundance of *E. coli*, *P. aeruginosa* as well as the total biomass under LDIR and 1% QAC (A, solid line) or LDIR and 1‰ QAC (B, solid line) as well as only 1% QAC (A, dotted line) or 1‰ QAC (B, dotted line).

**Figure S8:**
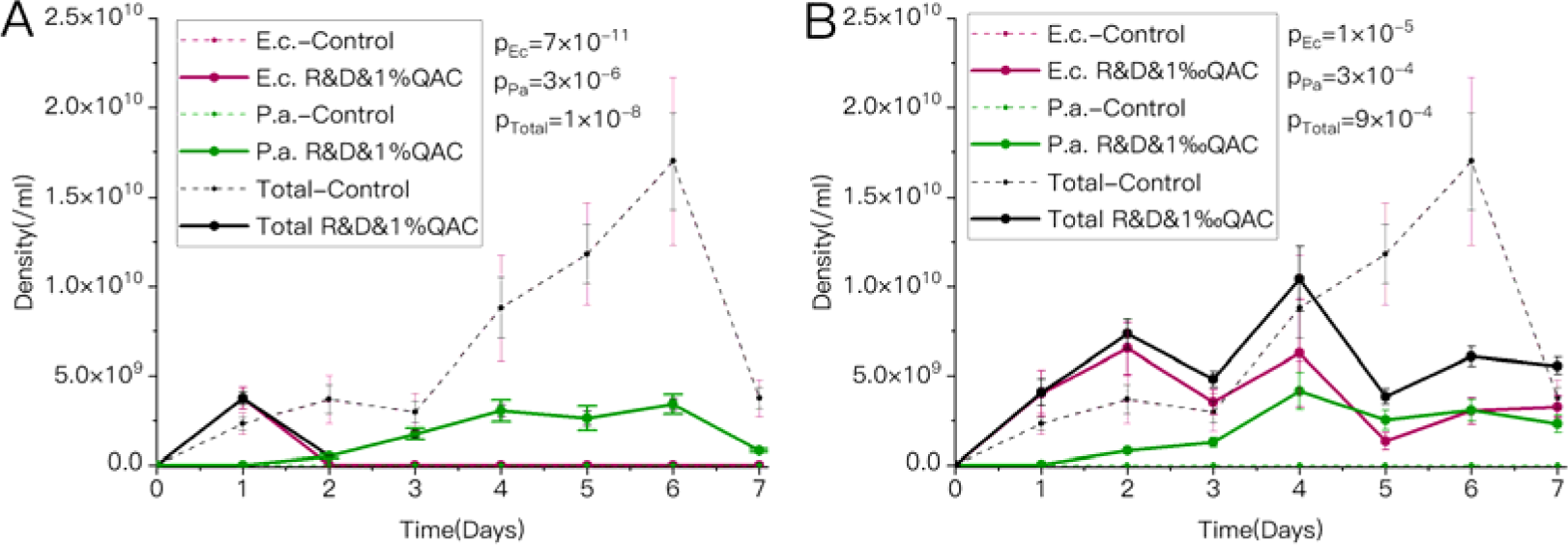
The absolute abundance of *E. coli*, *P. aeruginosa* as well as the total biomass under LDIR, 1% dilution and 1% QAC (A, solid line) or LDIR, 1% dilution and 1‰ QAC (B, solid line) as well as in the control group (dotted line).

**Figure S9:**
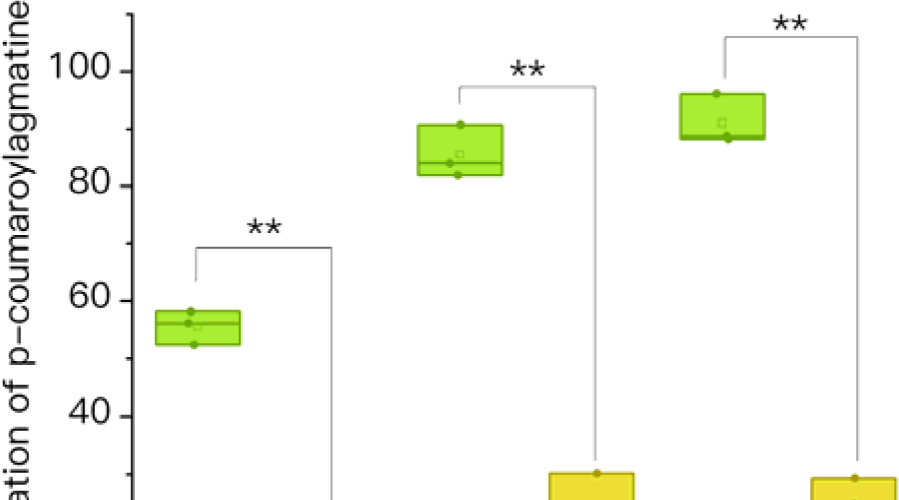
the concentration of p-coumaroyl agmatine in the group exposed under LDIR between the 4-6^th^ days dropped significantly compared with the control group. “**” meant (<).

**Figure S10:**
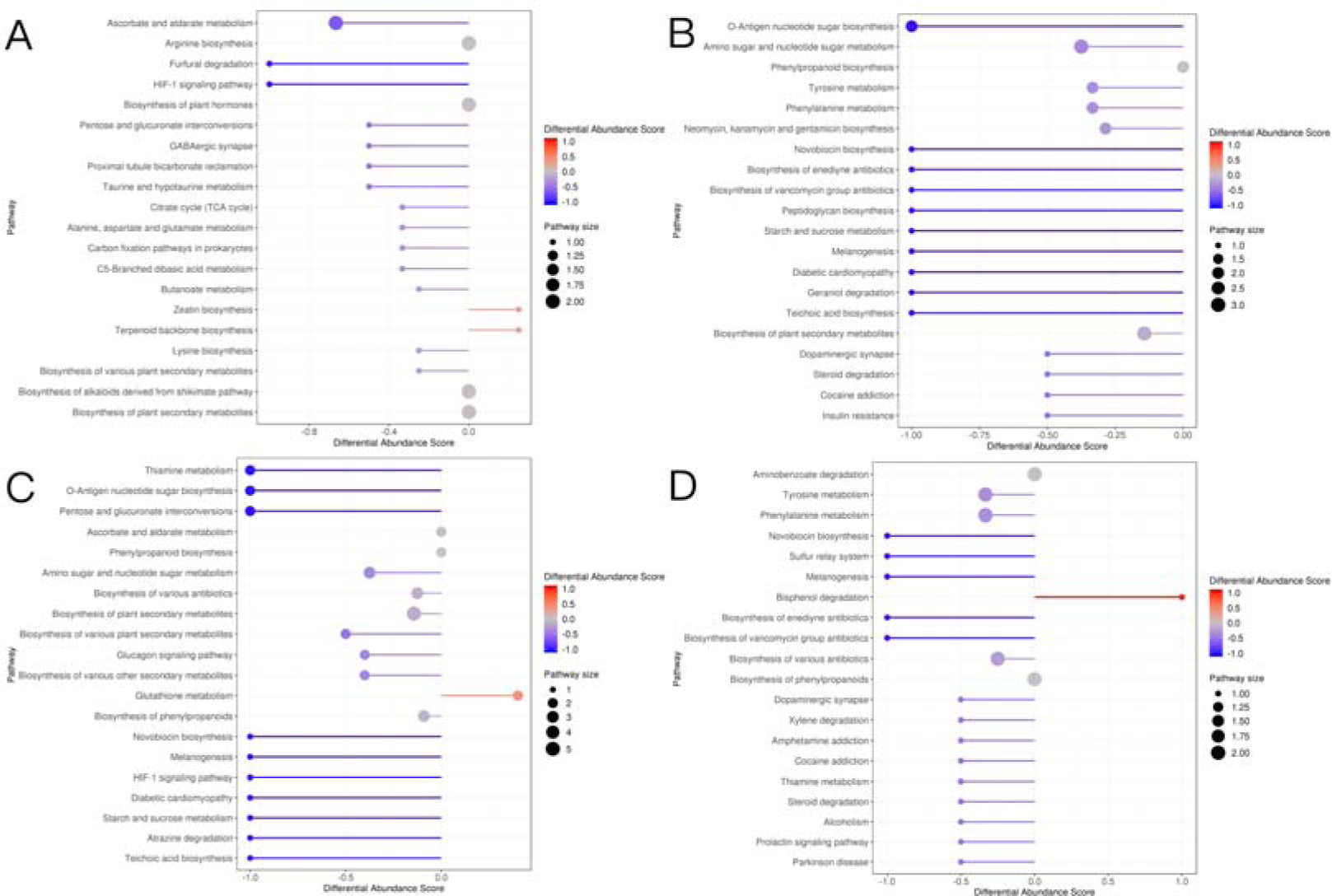
the differential abundance scores of differential metabolisms between the group under LDIR and the control group. A-D represent the 1, 4, 5, 6^th^ day, respectively.

**Figure S11:**
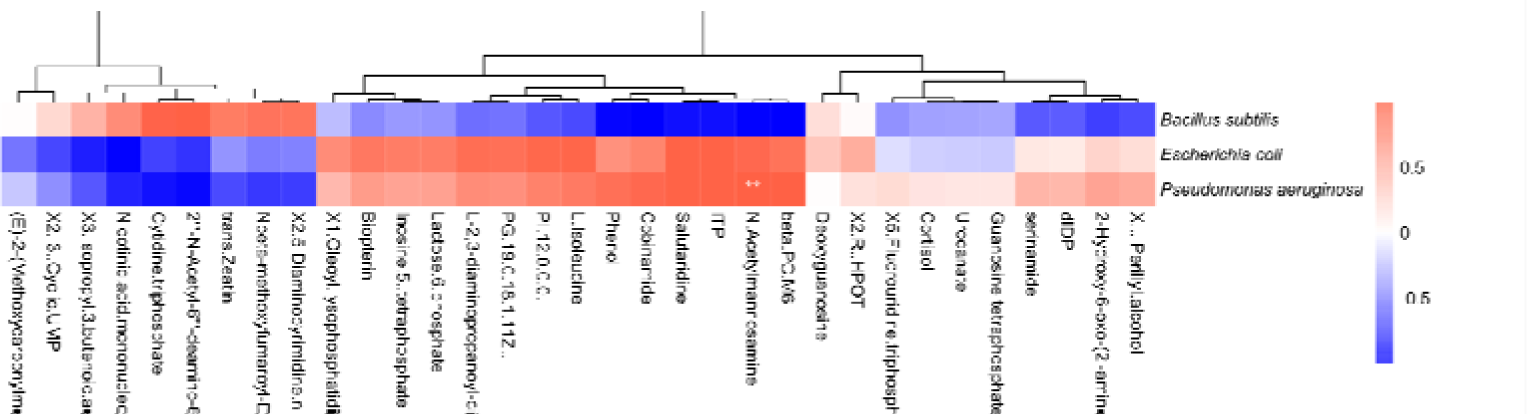
the enrichment of N-acetylmannosamine was markedly correlated with the rise of the abundance of *P. aeruginosa* (<) on the 4^th^ day of the group diluted with 1‰ ratio per day, compared with the control group. Metabolites with the top 10 highest concentrations in either group, or whose were considered in the intergroup correlation analysis. “**” meant (<), and “*” meant (<). The same is as below.

**Figure S12:**
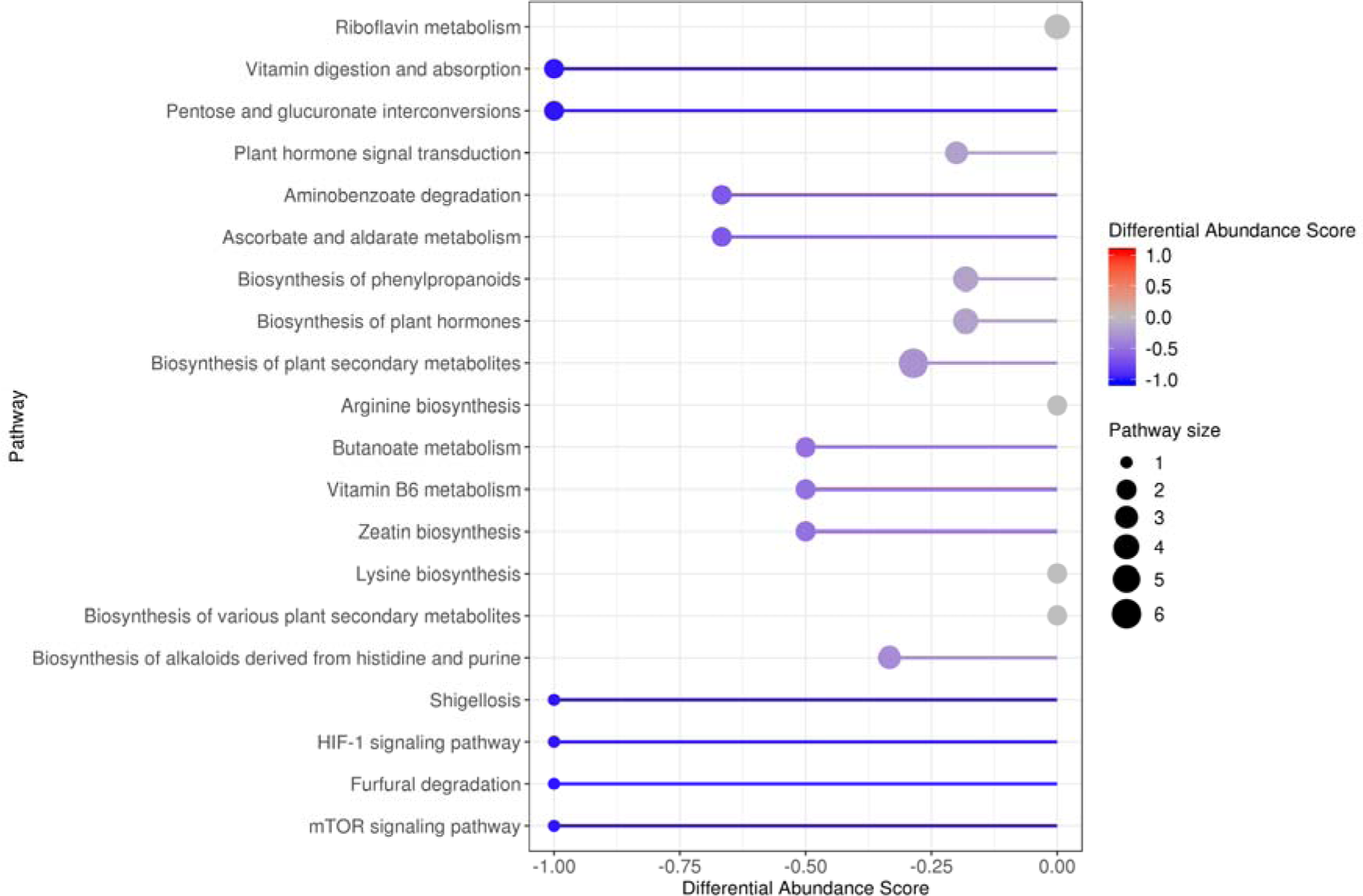
the differential abundance scores of differential metabolisms on 4^th^ day between the group diluted with 1‰ ratio per day and the control group.

**Figure S13:**
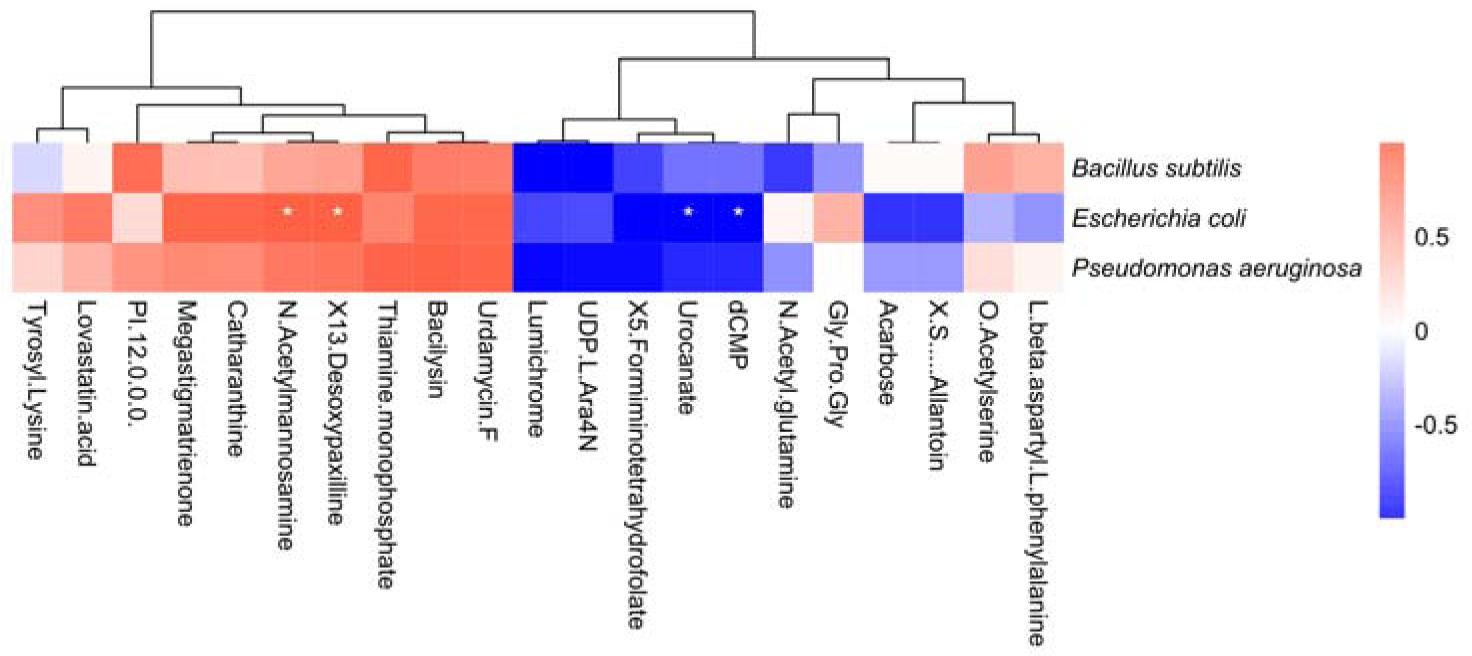
the enrichment of N-acetylmannosamine was markedly correlated with the rise of the abundance of *E. coli* (<) on the 4^th^ day of the group with 1% stock solution of QAC, compared with the control group.

**Figure S14:**
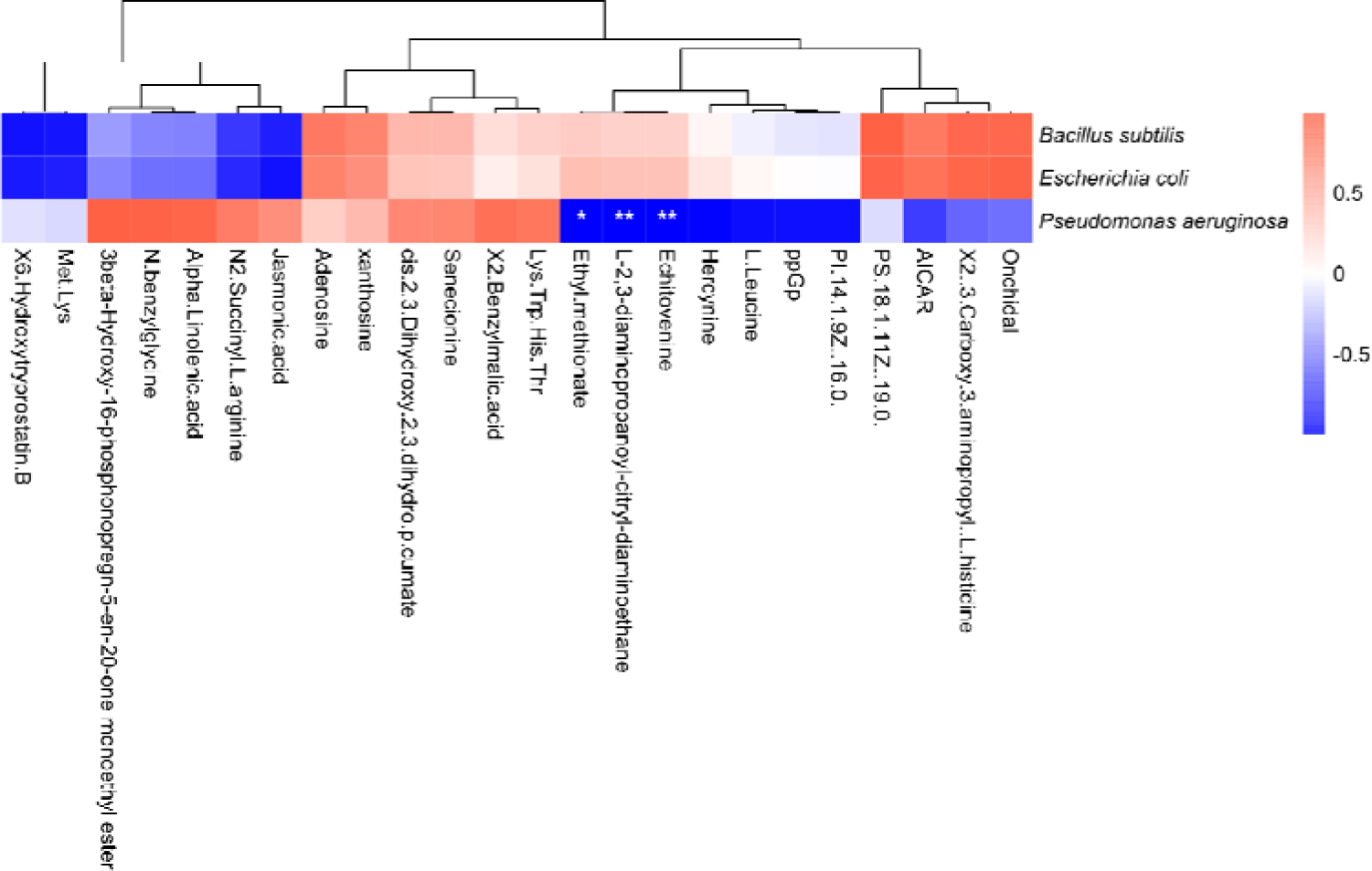
the drop of echitovenine was markedly correlated with the increase of the abundance of *P. aeruginosa* (<) on the 7^th^ day of the group with 1‰ stock solution of QAC, compared with the control group.

**Figure S15:**
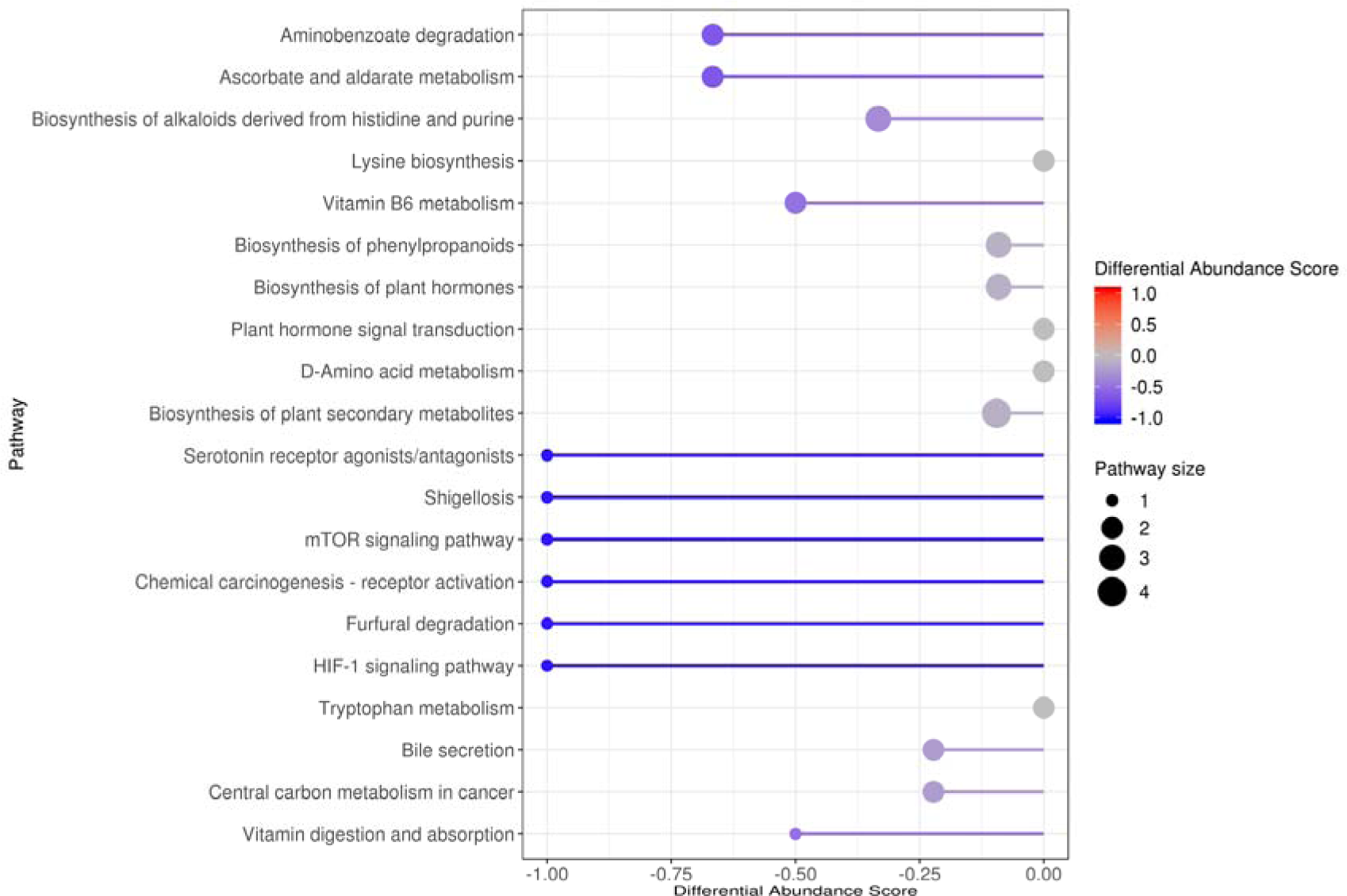
the differential abundance scores of differential metabolisms on 7^th^ day between the group with 1‰ stock solution of QAC and the control group.

**Figure S16:**
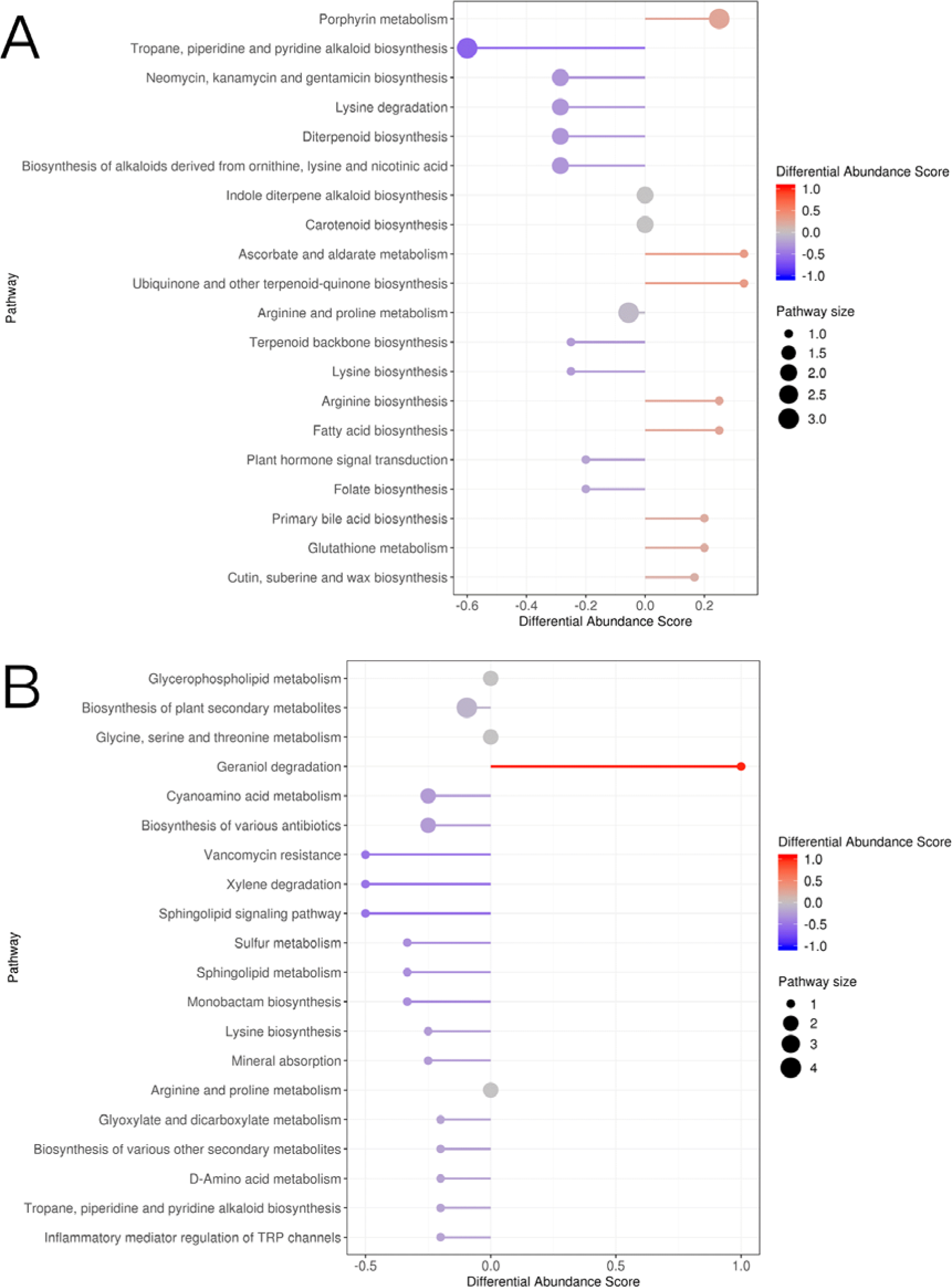
the differential abundance scores of differential metabolisms between the group with both LDIR and 1% stock solution of QAC and the group under 1% stock solution of QAC. A and B represent the 5^th^ and 6^th^ day, separately.

**Figure S17:**
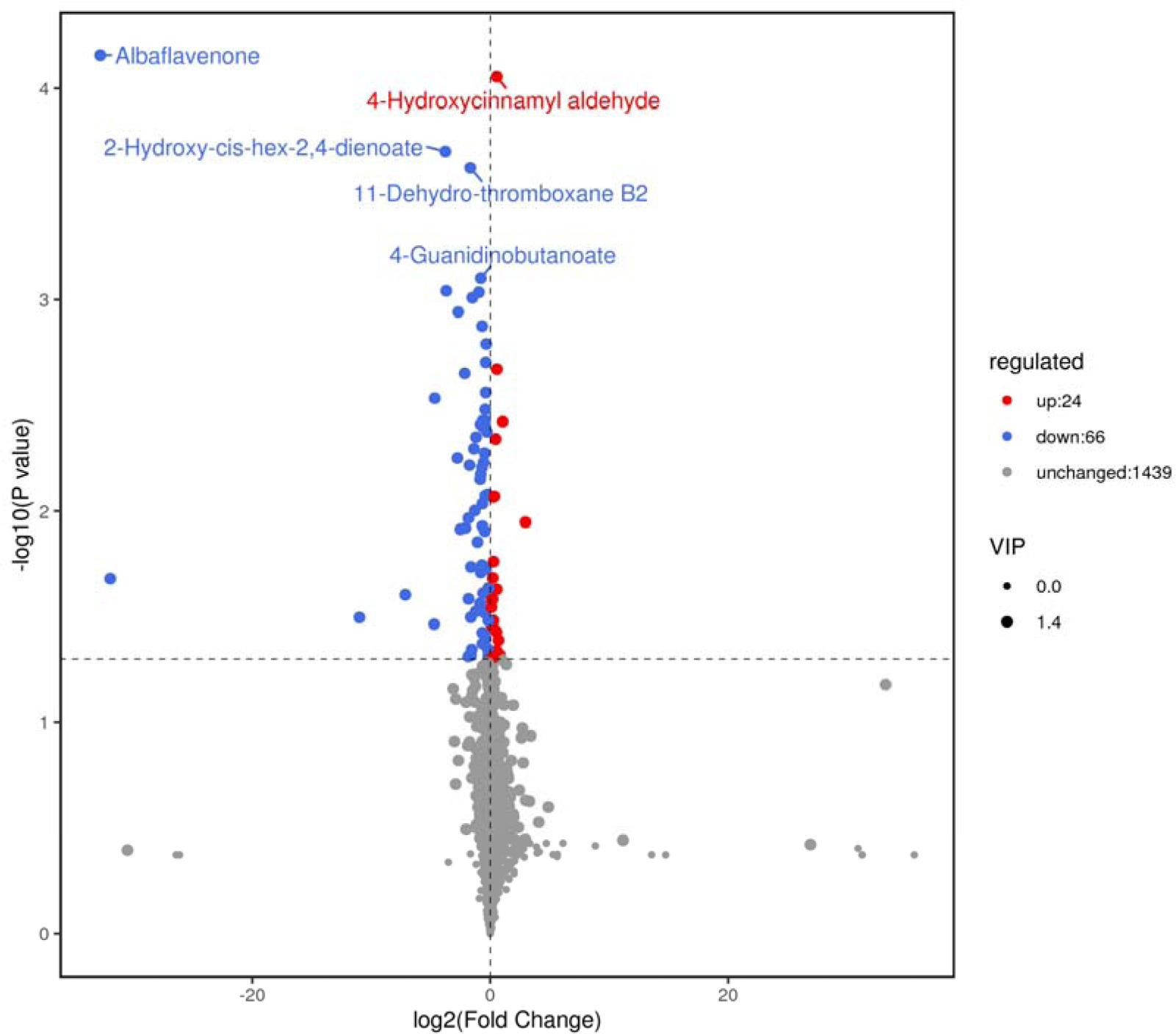
the concentration of albaflavenone sharply diminished on 4^th^ day in the group with both LDIR and 1‰ stock solution of QAC, compared with the control group.

**Figure S18:**
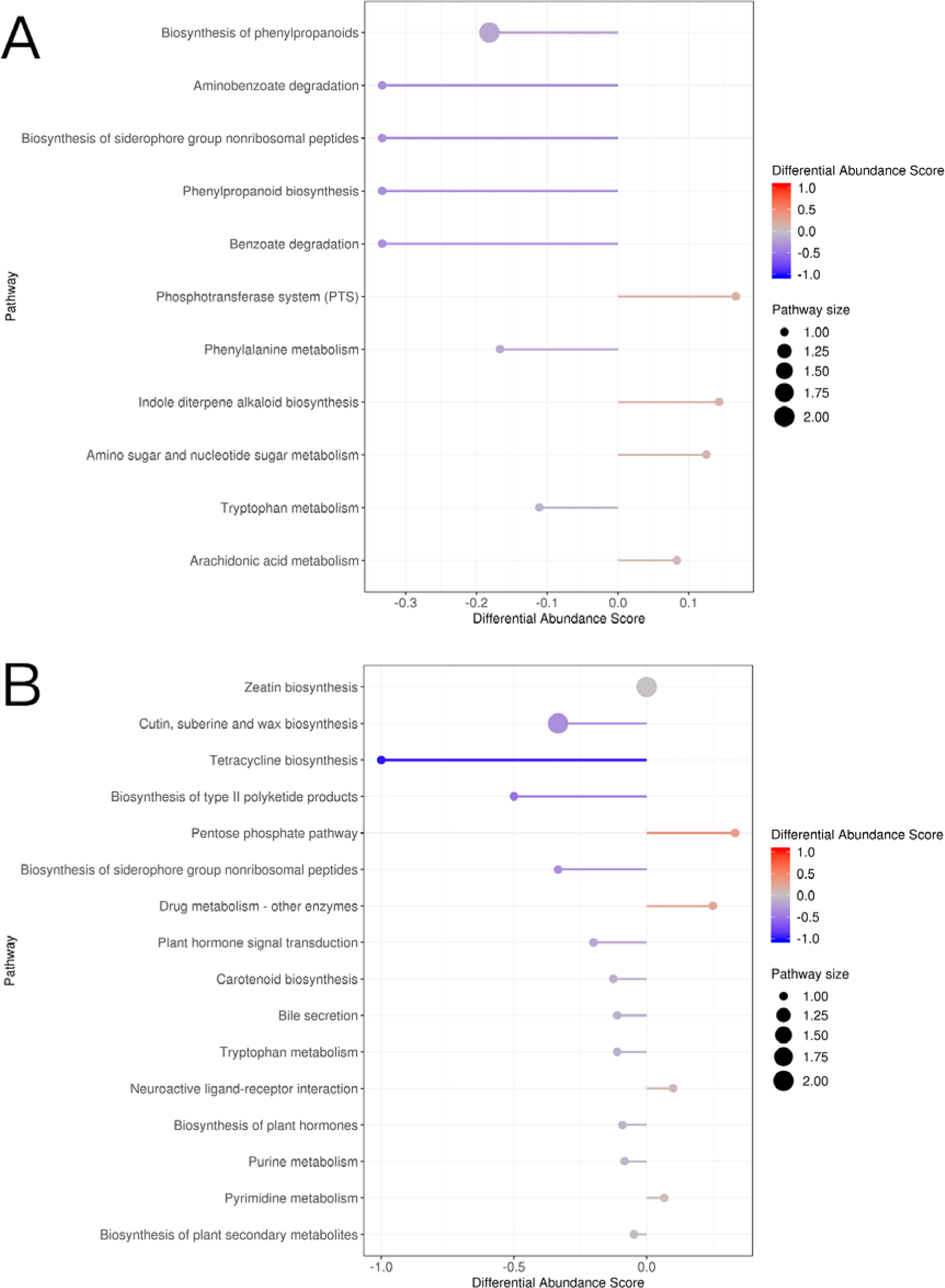
the differential abundance scores of differential metabolisms between the group with both LDIR and 1‰ stock solution of QAC and the group under 1‰ stock solution of QAC. A and B represent the 4^th^ and 7^th^ day, separately.

**Figure S19:**
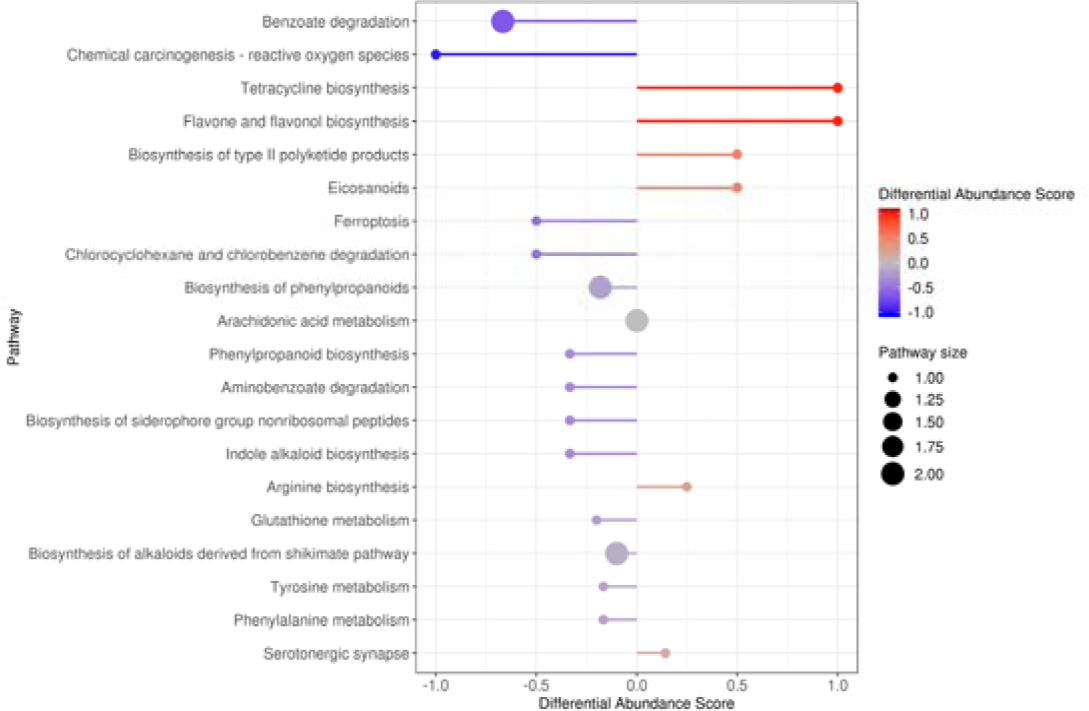
the differential abundance scores of differential metabolisms on 4^th^ day between the groups with 1% stock solution of QAC and 1‰ stock solution of QAC.

**Figure S20:**
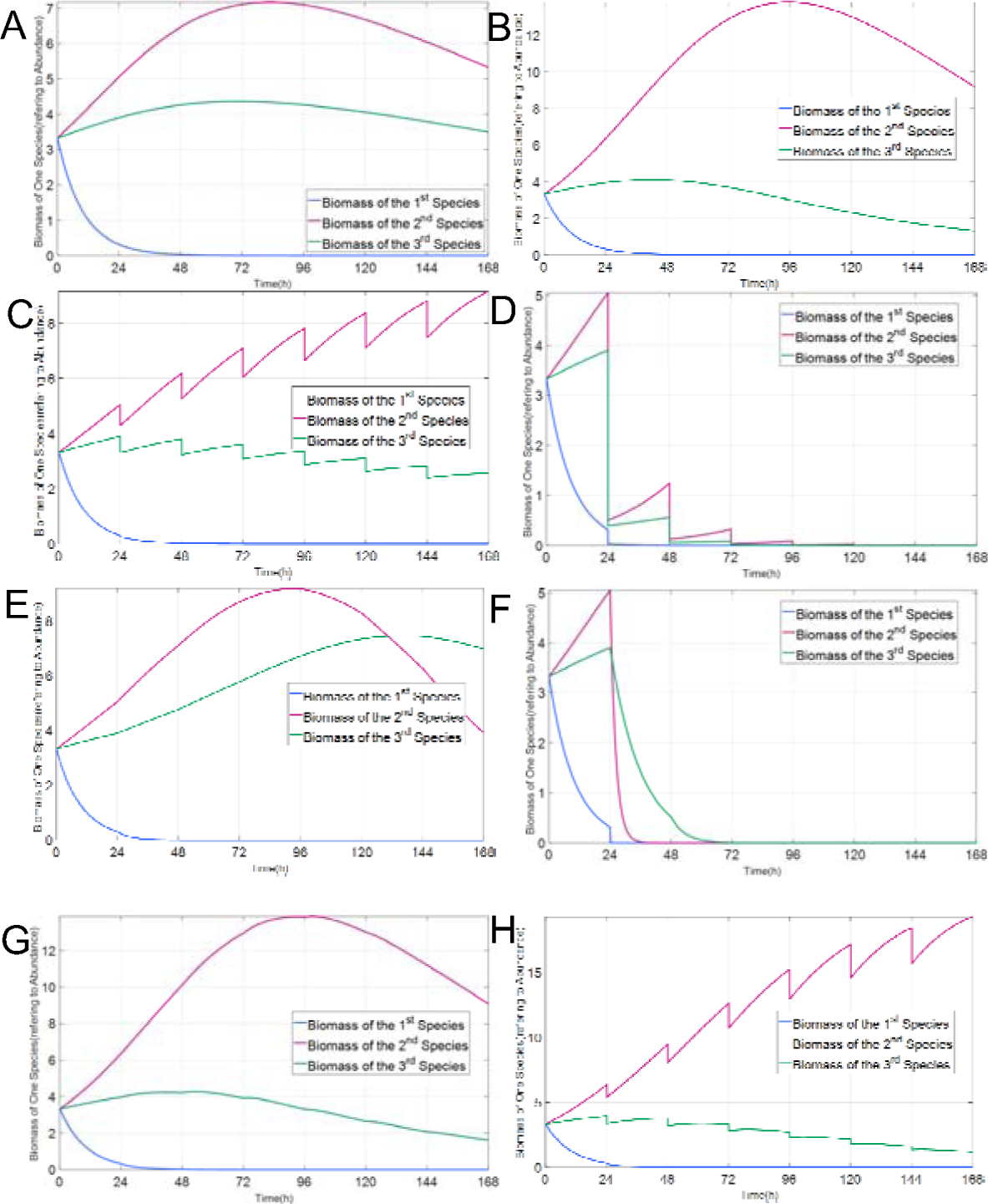
The simulation results from the model built in MatLab/Simulink. A-H represent the control group, LDIR, low dilution rate, high dilution rate, low QAC concentration, high QAC concentration, LDIR with low QAC concentration, all three factors together, respectively.

**Figure S21:**
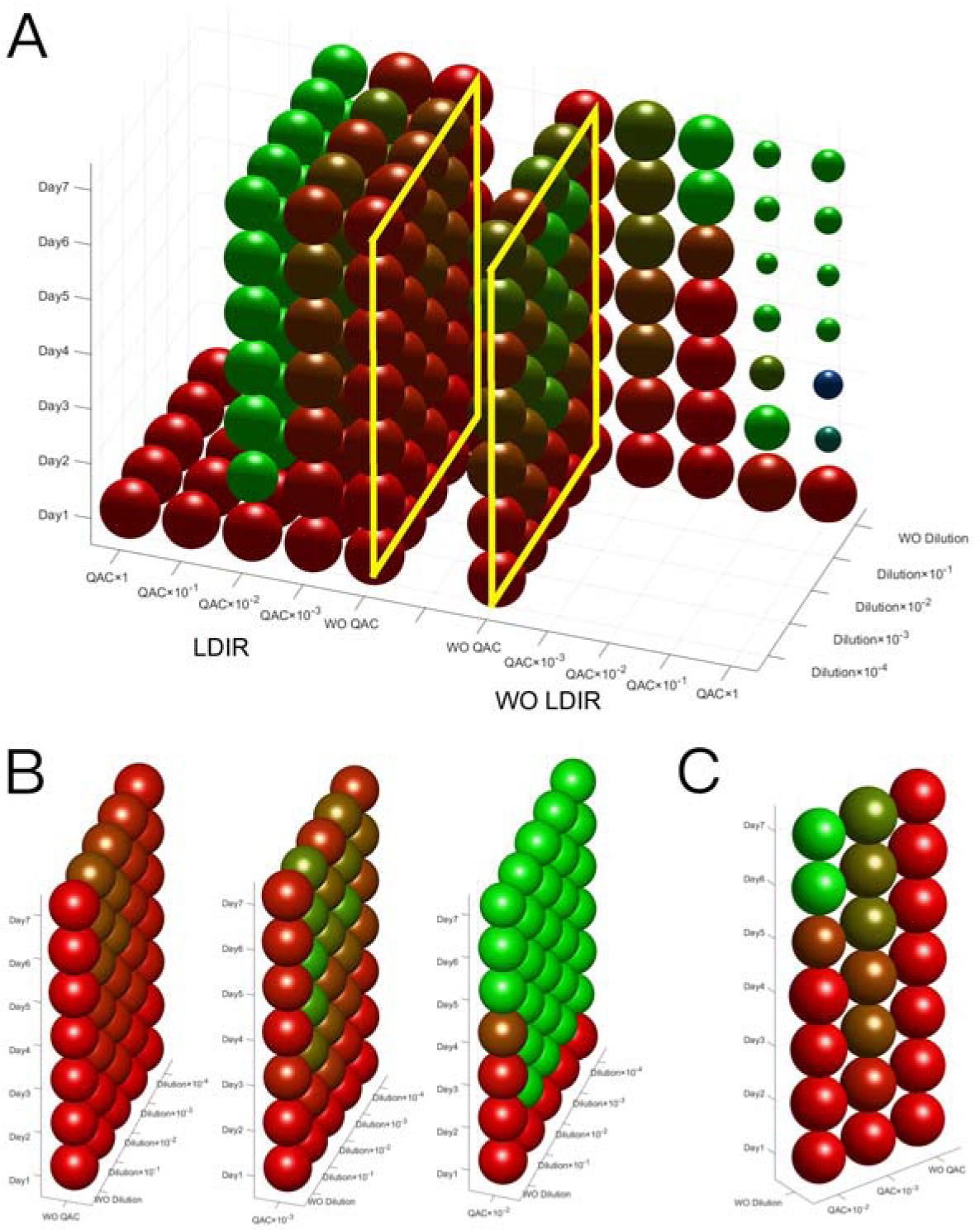
The overview of the community succession under all treatments and its slides. (A) The two planes in the yellow boxes showed that *E.coli* took the majority when both dilution and LDIR were applied compared to only dilution. (B) With higher gradients of QAC under 1%, *P. aeruginosa* gradually became the dominant species with LDIR. (C) With higher gradients of QAC under 1%, *P. aeruginosa* gradually became the dominant species without LDIR.

**Figure S22:**
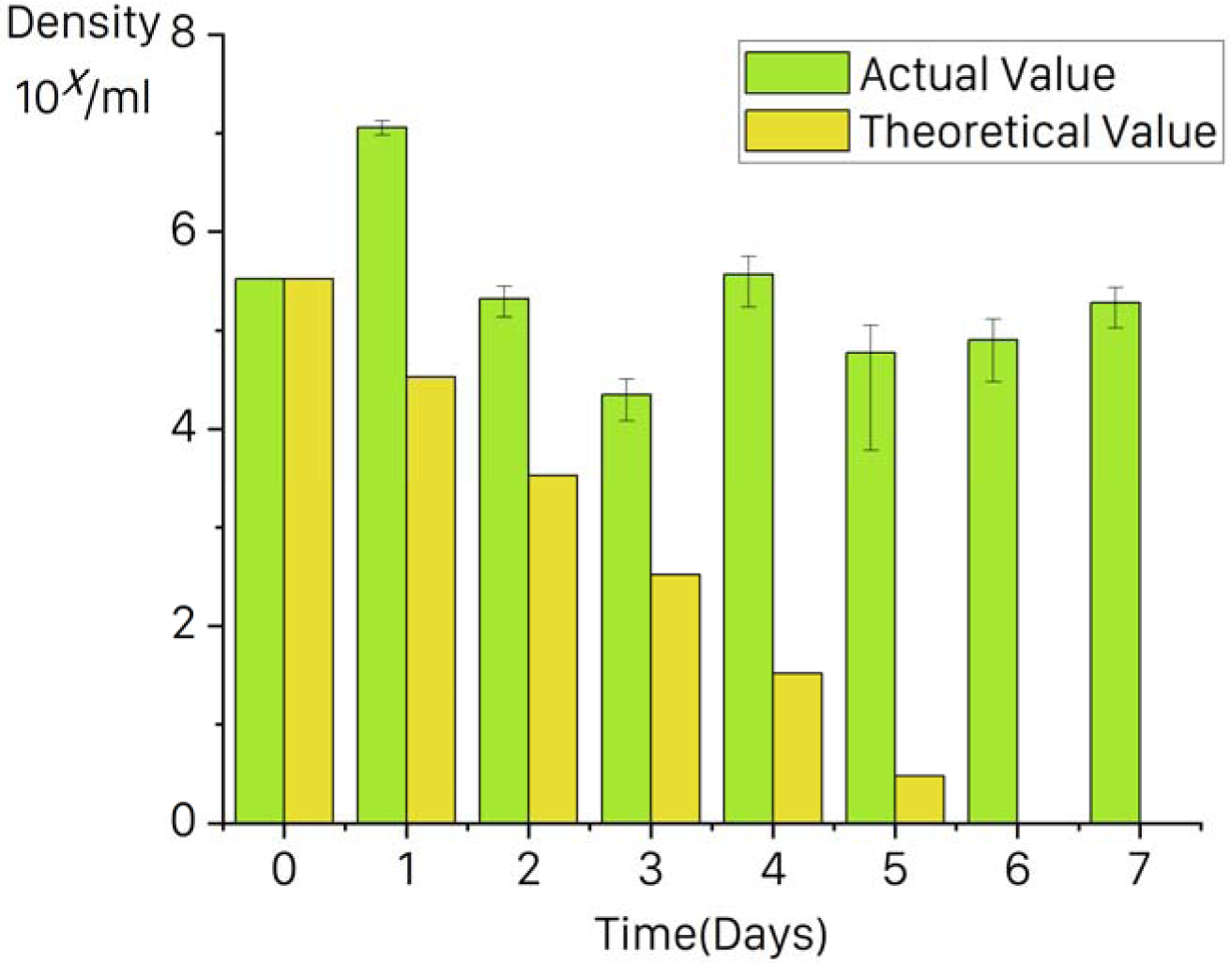
the actual and theoretical counts for 10× gradient dilution of *B. subtilis* cells.

**Table S1.** The composition of modified minimal R2A.

| Component | per 1000ml | Final concentration |
| --- | --- | --- |
| NaCl | 18.0g | 1.8% |
| Peptone | 0.5g | 0.05% |
| Yeast Extract Powder | 0.5g | 0.05% |
| Casein Peptone | 0.5g | 0.05% |
| Sodium Pyruvate | 0.3g | 0.03% |
| ddH<sub>2</sub>O | 1000ml |  |

**Table S2.** Conditions of Distinct Groups.

| Group | LDIR | Dilution | QAC |
| --- | --- | --- | --- |
| 1 | N | N | N |
| 2 | N | 10% | N |
| 3 | N | 1% | N |
| 4 | N | 1‰ | N |
| 5 | N | 0.1‰ | N |
| 6 | N | N | 100% |
| 7 | N | N | 10% |
| 8 | N | N | 1% |
| 9 | N | N | 1‰ |
| 10 | Y | N | N |
| 11 | Y | 10% | N |
| 12 | Y | 1% | N |
| 13 | Y | 1‰ | N |
| 14 | Y | 0.1‰ | N |
| 15 | Y | N | 100% |
| 16 | Y | 10% | 100% |
| 17 | Y | 1% | 100% |
| 18 | Y | 1‰ | 100% |
| 19 | Y | 0.1‰ | 100% |
| 20 | Y | N | 10% |
| 21 | Y | 10% | 10% |
| 22 | Y | 1% | 10% |
| 23 | Y | 1‰ | 10% |
| 24 | Y | 0.1‰ | 10% |
| 25 | Y | N | 1% |
| 26 | Y | 10% | 1% |
| 27 | Y | 1% | 1% |
| 28 | Y | 1‰ | 1% |
| 29 | Y | 0.1‰ | 1% |
| 30 | Y | N | 1‰ |
| 31 | Y | 10% | 1‰ |
| 32 | Y | 1% | 1‰ |
| 33 | Y | 1‰ | 1‰ |
| 34 | Y | 0.1‰ | 1‰ |

\*“N” means this condition was not applicable in this group. The ratio of dilution meant that part of the culture from $t$ -th day was taken into the R2A solution for gaining the 10ml new culture of $t + 1$ -th day. No dilution meant that R2A solution was added into the old culture to gain the 10ml new culture. The ratio of QAC meant the stock solution of QAC was added into the culture to make that concentration of the stock solution of QAC.

**Table S3.** The composition of QAC.

| Component | per 100ml | Final Concentration |
| --- | --- | --- |
| Octyl Decyl Dimethyl Ammonium Chloride (Purity: 80%) | 0.5g | 0.04% |
| Diocetyl Dimethyl Ammonium Chloride (Purity: 80%) | 0.25g | 0.02% |
| Didecyl Dimethyl Ammonium Chloride (Purity: 95%) | 0.21g | 0.02% |
| Alkyl Dimethyl Benzyl Ammonium Chloride, (50% C14, 40% C12, 10% C16) (Purity: 80%) | 0.66g | 0.0532% |
| ddH<sub>2</sub>O | 98ml | - |

**Table S4.**
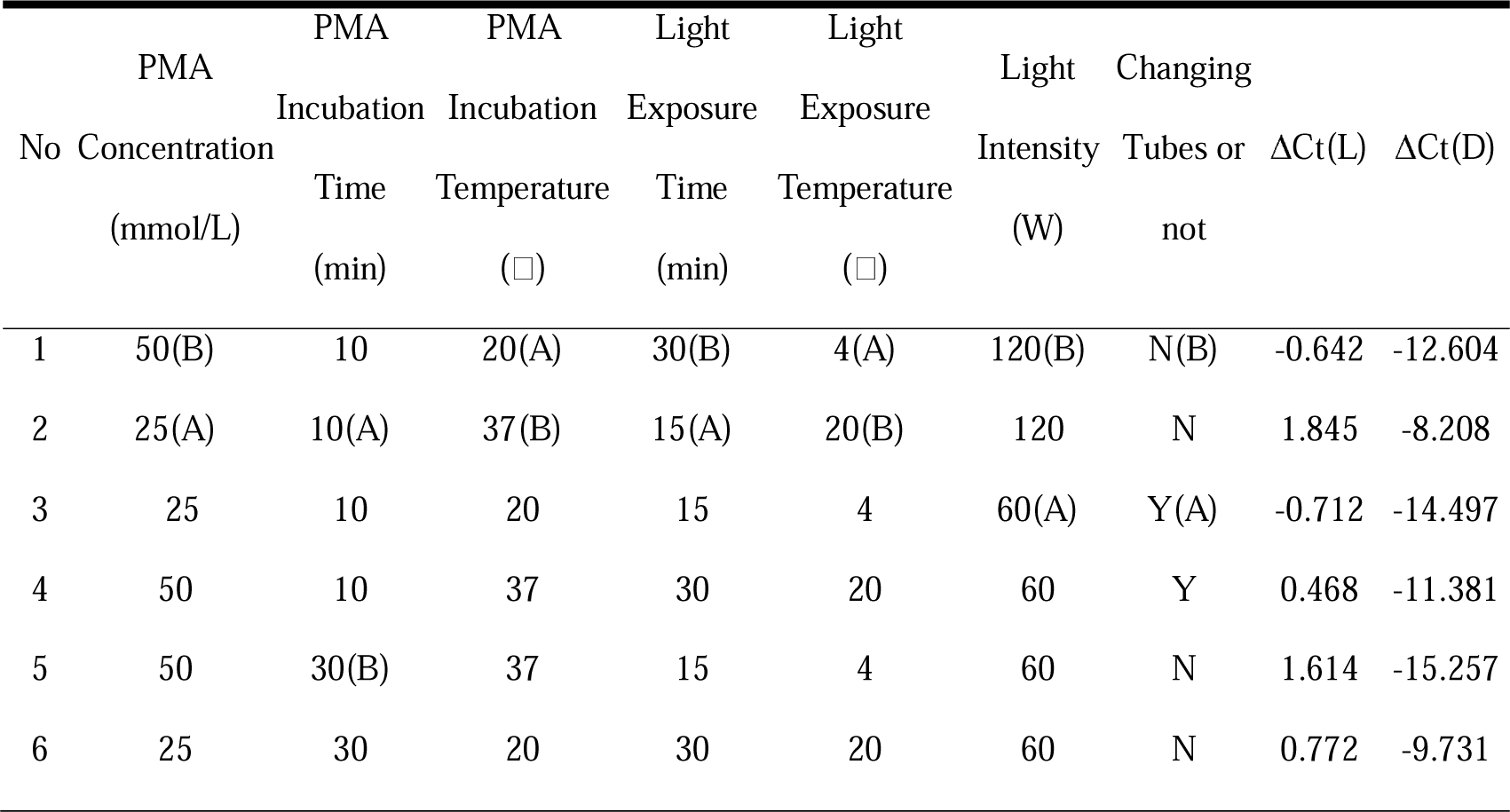
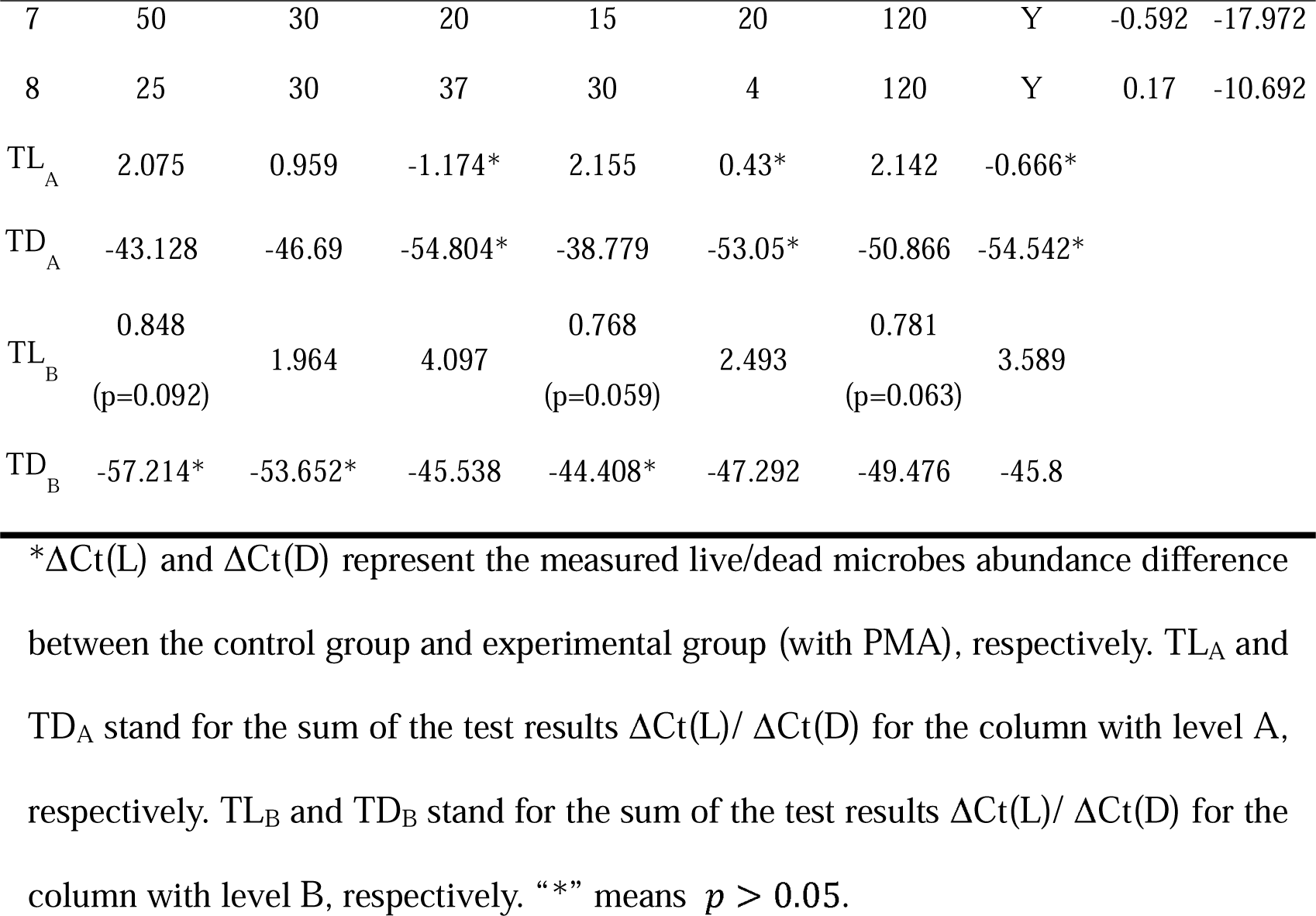
Orthogonal Experiments for PMA-qPCR Conditions.

**Table S5.**
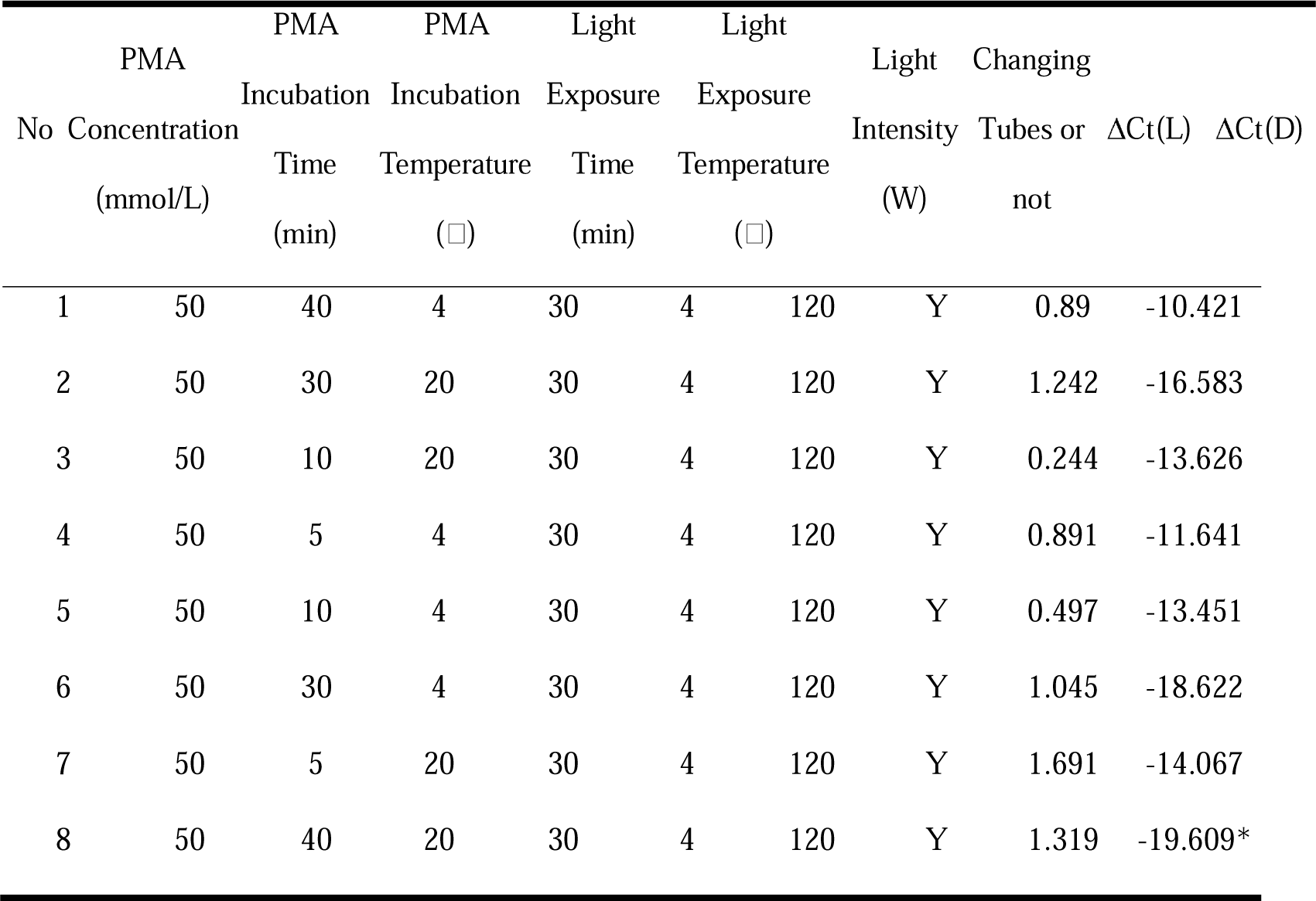
Gradient Experiments for PMA-qPCR Conditions.

## Supplementary Material A

Several studies from different perspectives have shown a special phenomenon in *Bacillus subtilis* (Kirchhoff & Cypionka, 2017; Tut et al., 2021; Xie et al., 2016), which is, that some nucleic acid dyes like PMA could penetrate a large number of living cells due to the excessively strong cell membrane potential. Therefore, at the macro level, the rate of false-negative qPCR results would increase, and the threshold for detection of living *Bacillus subtilis* cells through the PMA-qPCR method would spike. In the pre-experiments of this study, *Bacillus subtilis* did not lead in the community. Hence, daily dilution to 10% of the culture yesterday experiment was performed, and the magnitude of counts of *Bacillus subtilis* was much higher than the circumstance “No *Bacillus subtilis* survived, its DNA was diluted simultaneously”, as shown in Fig. S22. Therefore, direct qPCR was applicable for approximately counting *Bacillus subtilis*.

## Supplementary Material B

Eliminating the DNA of dead bacteria cells is the key aim of PMA in PMA-qPCR. The standard PMA protocol might be suitable for natural environmental samples, but not enough for high-abundance samples, like artificial microbial communities used in this study. Therefore, it is important to optimize its conditions. Related factors included PMA concentration, PMA incubation (with oscillation at 100rpm) time, PMA incubation temperature, light exposure time, light exposure temperature, light intensity (expressed as the blue LED bulb power), whether changing tubes between incubation and light exposure (Banihashemi et al., 2012; Baymiev et al., 2020; Codony et al., 2020; Emerson et al., 2017; Fittipaldi et al., 2012; Fleischmann et al., 2021; Loozen et al., 2011; Martin et al., 2013). To find the highest DNA of dead bacteria cells elimination rate, orthogonal experiments (Table S4) were implemented first on *P. aeruginosa* to screen the better condition of the two-level factor (PMA was not applicable for *B. subtilis* as seen in Sup. A and the primer of *E. coli* used for this study had a relatively high detection threshold). PMAxx was used as the improved reagent. Conditions were selected in previous studies (Agustí et al., 2018; Codony et al., 2020; Emerson et al., 2017; Fittipaldi et al., 2012; Fleischmann et al., 2021; García-Fontana et al., 2016; Kralik et al., 2010; Nocker et al., 2007).

The larger the absolute value of ΔCt(D), the higher the dead bacteria cells’ elimination rate. Based on no significant difference of ΔCt(D), the closer ΔCt(L) to zero, the smaller the influence of PMA on live cells. From the orthogonal experiments, these conditions were significantly better: the PMA concentration was 50 mmol/L, the light exposure time was 30min, the light exposure temperature was 4, and changing tubes were applied. 120W light intensity has a p-value of 0.063 so it was also selected.

Incubation time and temperature were tested another time in gradient experiments on *P. aeruginosa* as Table S5. As seen in Table, when the PMA incubation time was 40min and the PMA incubation temperature was 20, the absolute value of ΔCt(D) was the largest with a significant difference. Therefore, the optimal condition was: to incubate with 50mmol/L PMAxx (Cat. 40069, Biotium Inc., Hayward, CA, USA) 100rpm for 40min at 20, change a new tube, and expose it under a blue LED bulb still for 30min at 4.

When applying this optimal condition to *E. coli*, the ΔCt(D)=-18.268. Compared to the standard condition of PMA protocol, the detective elimination rate of dead *E. coli* surged from 25.976% to 99.999%, and that of *P. aeruginosa* dropped from 25.097% to 99.999%. The optimal condition can fulfill the need to measure high species abundance in this study and it was used when treating samples for these two species’ abundance data.

## Supplementary Material C

In this study, based on LC-QTOF, the qualitative and quantitative analysis of untargeted metabolomics of 72 samples was carried out. A total of 12,079 peaks were detected, of which 1,529 metabolites were annotated. The RSD ≤ 30% ones accounted for more than 60%, so it could be assumed that QC samples have good reproducibility, the instrument was stable, and the requirements of quality control were met.

In OPLS-DA (Orthogonal Projections to Latent Structures-Discriminant Analysis), all results of the experimental and their corresponding control groups could be completely separated, indicating significant overall changes in metabolites for each condition. In the comparison of groups J and V, the Q2Y < 0.5, thus VIP could not be considered as a screening condition for differential metabolites. Besides the above, Q2Y of all other comparisons > 0.5, indicating that the OPLS-DA model was valid. Permutation tests were performed to check the reliability of the OPLS-DA model. The slopes of Q2Y were positive for all groups except for the comparison of J and V, further supporting that the OPLS-DA model was significant. Most of the blue dots in the scatterplots were located above the red dots, indicating good independence of the OPLS-DA model.

In the following analysis, metabolites with the same trend on consecutive days, |*log*_2_*FC*| ≥ 20 (except for *SD* = 0 to remove the systematic error of the instrument) or ranking top 10 were screened.

